# Butyrophilin 2A2 promotes T cell immunoregulation by enhancing CD45 phosphatase activity within the immune synapse

**DOI:** 10.1101/2024.11.12.623303

**Authors:** Shafat Ali, Anders H. Berg, Michifumi Yamashita, Ambart E. Covarrubias, Ruan Zhang, Vincent Dupont, Bong-Ha Shin, Shen Yang, Ramachandran Murali, Madhusudhanarao Katiki, Margareta D. Pisarska, Ravi Thadhani, Peter S. Heeger, Stanley C. Jordan, S. Ananth Karumanchi

## Abstract

B7 costimulatory family member Butyrophilin 2A2 (BTN2A2) is predominantly expressed by antigen presenting cells and regulates T cell immunity, but molecular mechanisms are unclear. Using immunoblots analyzing TCR-initiated signaling intermediaries, co-immunoprecipitation studies, confocal microscopy, structural modeling-guided mutational analyses, and microscale thermophoresis, we demonstrate that BTN2A2 directly interacts with CD45RO, resulting in CD45 retention within the immune synapse during TCR activation. Recombinant BTN2A2 increased murine CD4+Foxp3+ regulatory T cells (Treg) and reduced T helper 17 (Th17) cells in vitro through mechanisms dependent on CD45 phosphatase activity. BTN2A2 treatment reduced clinical expression of two murine autoimmune disease models and increased Treg/Th17 ratios. Analyses of BTN2A2-deficient animals showed exacerbated disease associated with reduced Treg/Th17 ratios. Addition of BTN2A2 to human mixed lymphocyte responses similarly enhanced human Treg and suppressed Th17 cells and was CD45 phosphatase dependent. Together, our studies identify BTN2A2 as a physiological CD45RO ligand that enhances CD45 phosphatase activity in murine and human T cells, providing mechanisms for BTN2A2-mediated amelioration of autoimmunity.

**Summary:** Butyrophilin 2A2 ameliorates autoimmunity by binding to CD45RO on activated T cell surfaces leading to dampened TCR signaling which in turn leads to expansion of T regulatory cells and reduction of Th17 differentiation.

## INTRODUCTION

T cells recognize antigens presented through peptide:MHC complexes using surface expressed, heterodimeric αβT cell receptors (TCRs). As the TCR lacks intrinsic kinase activity, signals initiated by TCR ligation involve recruitment of the Src family kinase Lck, which then phosphorylates immunoreceptor tyrosine-kinase-based motifs (ITAMs) within the TCR-associated CD3 ζ-chain. Lck also phosphorylates subsequently recruited Zap70 kinase, thereby propagating requisite downstream signals required for full T cell activation. Studies performed over the past 30 years showed T cell activation is controlled in part by cell surface expressed CD45 (Courtney et al., 2018).

CD45 is a transmembrane glycoprotein that contains an intracellular tyrosine phosphatase domain capable of dephosphorylating multiple TCR immunoreceptor tyrosine activation (ITAM) motifs. Differential splicing results in expression of multiple CD45 isoforms (i.e. RA, RB, RC, RO) (Hermiston et al., 2003). Following TCR stimulation, CD45 is initially recruited to the supramolecular activation cluster (SMAC) but is then expelled, segregating it from the TCR (Jung et al., 2021; Leupin et al., 2000). Evidence suggests that this segregation of CD45’s phosphatase activity from the TCR is essential for Lck-initiated signal propagation that results in full T-cell activation.

Conversely, retention of CD45 within the T cell immune synapse (IS) regulates the strength and duration of TCR activation (He et al., 2002; Hermiston et al., 2003), and perturbations of CD45 activity contribute to development of autoimmune disease (Majeti et al., 2000; Vang et al., 2008). Despite decades of work by multiple groups delineating these molecular mechanisms, it is not known how CD45 segregation versus retention during TCR activation may be regulated, or if ligands co-presented by antigen-presenting cells (APCs) play a role.

Butyrophilins (BTNs) are glycoproteins originally isolated from breast milk that have imprecisely understood immune-regulatory effects and are implicated in maintaining maternal-fetal tolerance (Ogg et al., 2004; Rhodes et al., 2016; Zhao et al., 2020). The mRNAs encoding for BTN and BTN-like molecules are widely expressed in lymphoid and non-lymphoid tissues (Abeler-Dorner et al., 2012). Butyrophilin immunoglobulin domains exhibit structural similarities to the B7 family of co-receptors, including B7- 1/CD80, B7-2/CD86, ICOS-L and PD-L1, and Butyrophilin 2A2 (BTN2A2) was previously shown to be expressed by professional APCs including B cells, macrophages, and dendritic cells (DCs)(Smith et al., 2010). Results of in vitro studies suggest that BTN2A2 can modulate T cell receptor (TCR) signaling and promote *de novo* Foxp3 expression (Ammann et al., 2013; Smith et al., 2010). Mice genetically deficient in BTN2A2 exhibit impaired CD4+ regulatory T cell function, potentiated anti-tumor immunity, and augmented clinical manifestations of experimental autoimmune encephalomyelitis, all of which were attributable to deficiency of BTN2A2 in APCs (Sarter et al., 2016). While these cumulative findings implicate a key immunoregulatory function for BTN2A2, the exact molecular mechanisms underlying these effects remain unclear.

Herein we demonstrate that BTN2A2 functions as a ligand for CD45, binding both to the TCR complex and CD45RO isoform on T cell surfaces, resulting in retention of CD45 phosphatase activity proximal to the TCR complex. Consequently, the enhanced phosphatase activity reduces downstream TCR signaling, which in turn promotes regulatory T cell (Treg) expansion, suppressing T effector cell differentiation. We further demonstrate the significant immunomodulatory effects of BTN2A2 in a mouse model of autoimmune glomerulonephritis, showing that BTN2A2 genetic deficiency exacerbated kidney injury, whereas administration of recombinant BTN2A2 limited disease severity. Furthermore, in a second mouse model of immune-mediated spontaneous abortion, treatment of pregnant dams with recombinant BTN2A2 helped animals tolerate their pregnancies, reducing fetal loss.

## RESULTS

### BTN2A2 blocks CD3-dependent signaling in Jurkat Cells

To elucidate the mechanisms underlying BTN2A2’s ability to regulate T cell immunity (Ammann et al., 2013; Sarter et al., 2016), we generated a soluble recombinant human BTN2A2-Fc fusion protein using baculoviral expression systems. Consistent with prior reports performed in murine systems (Ammann et al., 2013), recombinant human BTN2A2-Fc, but not recombinant Fc control protein, blocked anti-CD3-induced IL-2 production by Jurkat cells (Supplementary Fig. 1A-C).

When we next evaluated signaling components downstream of the TCR, we observed that BTN2A2-Fc reduced anti-CD3 induced phosphorylation of Zap70 and CD3ζ in Jurkat cells (Supplementary Fig. 2A). Addition of the phosphatase inhibitor pervanadate resulted in hyperphosphorylation of Zap70 and CD3ζ at baseline (unstimulated), and in contrast to the findings in the absence of pervanadate, addition of BTN2A2-Fc to the culture had no effect on the hyperphosphorylated proteins (Supplementary Fig. 2B).

Together these data suggested that BTN2A2-Fc activates a phosphatase that downregulates TCR signaling.

### BTN2A2 blocks TCR signaling by binding to and enhancing CD45 phosphatase

Based on the known molecular mechanisms linking CD45’s phosphatase activity to TCR signaling (Jung et al., 2021; Leupin et al., 2000), we tested the hypothesis BTN2A2’s inhibitory effect on TCR activation is mediated through interaction with CD45. Western blotting of CD45-associated proteins after co-immunoprecipitation (co-IP) of CD45 in unstimulated Jurkat cells demonstrated that in the absence of TCR activation, CD45 was normally associated with several components of the TCR complex (Zap70 and CD3ζ). However, activation of the TCR complex with anti-CD3 antibodies resulted in CD45 dissociating from Zap 70 and CD3ζ. In contrast, TCR activation in the presence of BTN2A2 caused CD45 to remain associated with the TCR complex (Fig. 1A).

**Figure 1:**
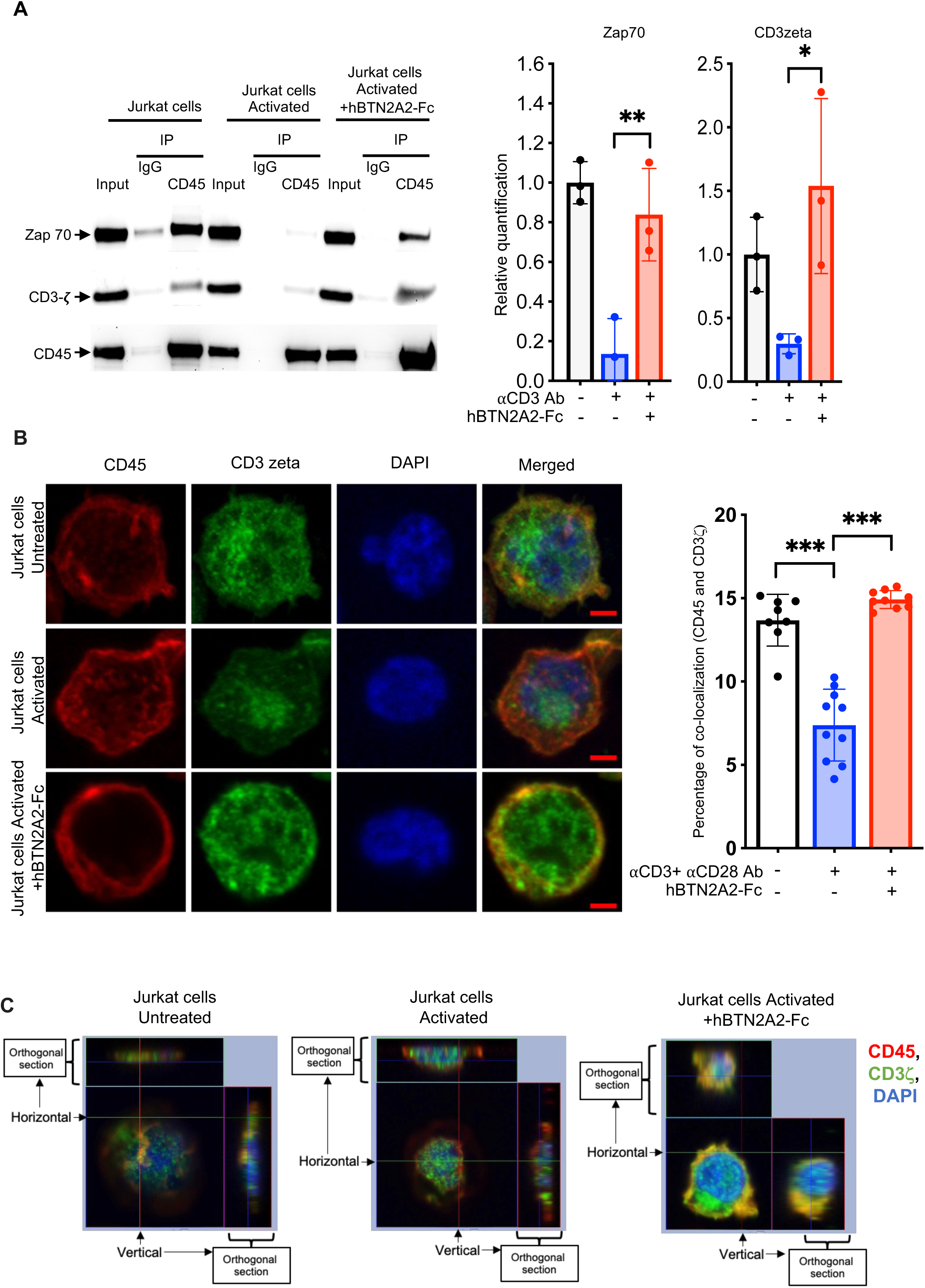
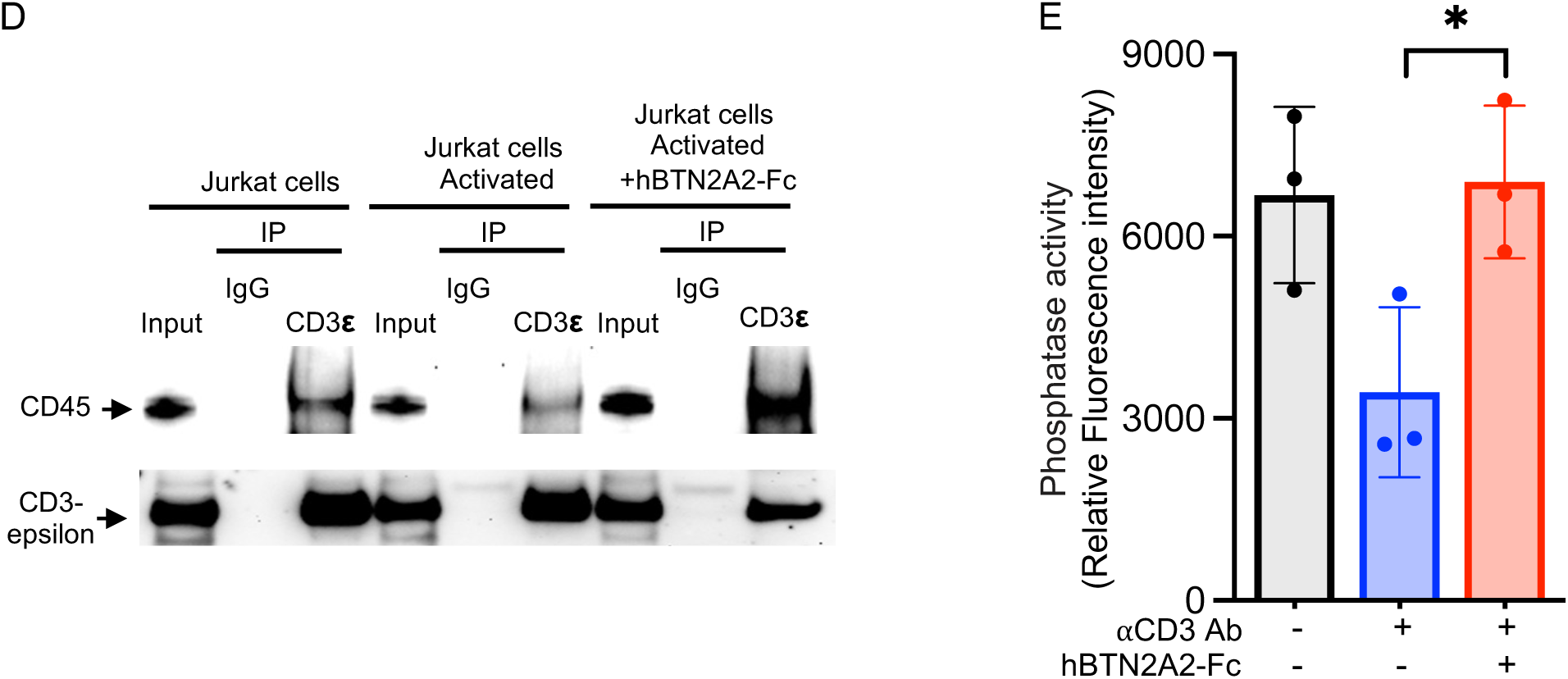
BTN2A2-Fc enhances interaction and co-localization of CD45 with TCR signaling proteins. (A) Jurkat cells were stimulated for 3 min with immobilized anti-CD3 antibody (10 μg/ml) in the presence or absence of BTN2A2-Fc (10 μg/ml). Cells were lysed in IP buffer, immunoprecipitated with anti-CD45 antibody and immunoblotted for total Zap70 and CD3*ζ*. Input is 5% of cell lysate. Right panel show intensity plots depicted as mean±SD (N=3) (B) Immunostaining and confocal microscopy analysis shows segregation of CD45 from CD3*ζ* in Jurkat cells treated with or without immobilized BTNA2-Fc protein (10 μg/ml) and activation by plate-bound anti-CD3/anti-CD28 antibody (10 μg/ml each) for 3 min. Right panel shows quantification of CD3*ζ* and CD45 colocalization in the presence or absence of BTN2A2-Fc proteins from multiple fields. More than 500 cells from each group were included in the analysis shown in the right panel. Magnification 63x resolution with Zeiss ‘Immersol’ immersion oil (refractive index of 1.518) (C) High resolution immunostaining analysis showing single cell orthogonal sections (z-stacks) in Jurkat cells with the same condition as in Figure 1B. (D) Co-immunoprecipitation experiments were performed using anti-CD3ε antibody in Jurkat cells with the same condition as in (A), followed by immunoblot with anti-CD45 antibody. Input is 2% of cell lysate. (E) CD45-specific phosphatase activity measured in immunoprecipitate from samples in panel C using FDP (Fluorescein Diphosphate, Tetraammonium Salt) substrate as described in the methods. Data depicted as mean ± SD (N=3). One-way ANOVA with Tukey’s multiple comparison test; *p*<0.05 (*), *p*<0.01 (**), *p*<0.001 (***).

Similarly, when we performed co-IP experiments with Zap70 pulldown in activated Jurkat cells treated with BTN2A2, CD45 associated with Zap70 (Supplemental Fig. 3).

Cellular imaging with confocal microscopy confirmed colocalization of CD45 and CD3ζ in untreated cells at baseline, and segregation of CD45 following anti-CD3/anti- CD28-induced TCR activation. CD3ζ staining following TCR activation was largely perinuclear as previously noted (Bunnell et al., 2002). Interestingly, we noted maintenance of CD45 and CD3ζ colocalization when cells were activated in the presence of BTN2A2, suggesting that BTN2A2 prevents the exclusion of CD45 from the immune synapse during the early phases of T cell activation. (Fig. 1B-C).

Co-IP experiments additionally demonstrated that compared to unstimulated cells, anti- CD3 activation reduced the association of CD3ε with CD45, whereas TCR activation in the presence of BTN2A2 augmented the interaction between CD45 and CD3ε (Fig. 1D). Functional experiments looking at TCR-associated CD45 phosphatase activity showed that when TCR complexes were isolated by co-IP with anti-CD3ε antibodies, CD45 phosphatase activity was detectable at baseline, decreased following TCR activation, and maintained when cells were activated in the presence of BTN2A2, suggesting segregation of CD45 from the TCR complex after activation and retention in the presence of BTN2A2 (Jung et al., 2021; Leupin et al., 2000) (Fig. 1E). Together with prior studies that activating CD45 phosphatase on lipid microdomains on T cell surfaces results in decreased sensitivity of TCR-mediated signaling (He et al., 2002), our new data indicate BTN2A2 dampens TCR signaling by activating CD45 phosphatase activity in the TCR complex which in turn blocks critical phosphorylation events downstream of the TCR.

### BTN2A2 interacts with CD45 phosphatase

We next employed a co-IP experimental strategy using Jurkat cells to test for direct interactions between CD45 and endogenously expressed BTN2A2 (Fig. 2A). These assays showed CD45 co-immunoprecipitated with endogenous BTN2A2 in unstimulated Jurkat cells, and binding was significantly enhanced in activated Jurkat cells. Anti-CD3 activation of Jurkat cells did not change endogenous CD45 and BTN2A2 expression levels (Supplementary Fig. 4A). Complementary experiments with exogenous BTN2A2- Fc or Fc protein control confirmed that exogenous BTN2A2-Fc but not Fc-tag protein binds to CD45 in activated T cells (Fig. 2B-C). This interaction between CD45 and BTN2A2 in activated T cells was specific as other abundantly expressed cell surface proteins such as CD43 failed to interact with BTN2A2 under similar conditions as above (Supplemental Fig. 4B).

**Figure 2:**
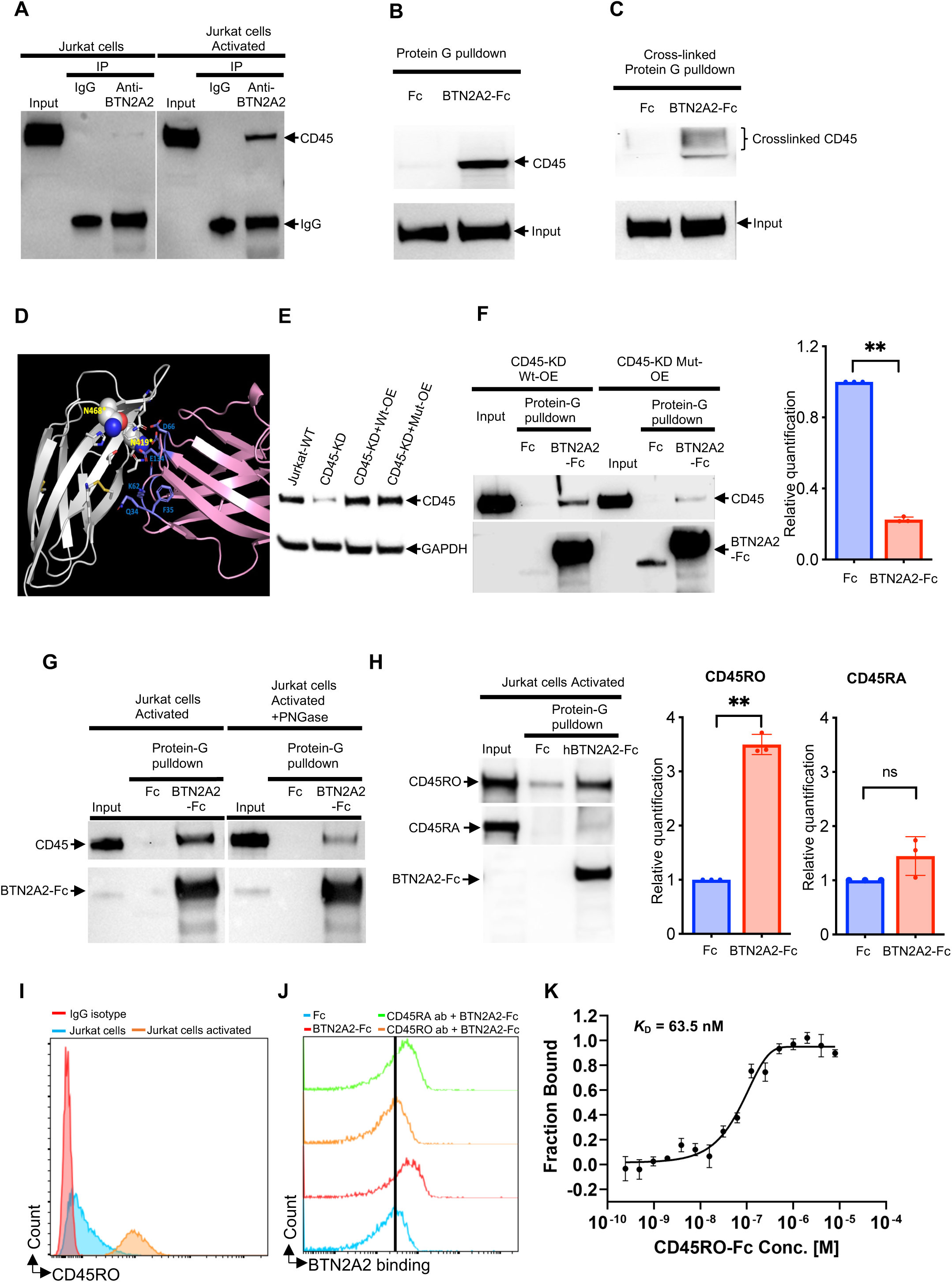
CD45 phosphatase co-immunoprecipitated with BTN2A2. (A) Jurkat cells were immunoprecipitated with anti-BTN2A2 antibody or IgG and blotted with CD45 antibody at unstimulated state or after activation using anti- CD3 and anti-CD28 antibody (1 μg/ml each) for 48 hours. Input is ∼2% of cell lysate. One representative experiment is depicted. (B) Jurkat cells activated with anti-CD3 antibody and anti-CD28 antibody (1μg/ml each) for 48 hours were incubated with BTN2A2-Fc or Fc-tag protein as described in Methods. Following Protein-G pulldown, eluted proteins were immunoblotted with anti-CD45 antibody or BTN2A2 antibody. Input is ∼2% of cell lysate. One representative experiment is depicted. (C) Jurkat cells activated with anti-CD3 antibody and anti-CD28 antibody (1μg/ml each) for 48 hours were incubated with recombinant BTN2A2-Fc or Fc-tag protein and crosslinked with BS3 as described in the methods. Cell lysate was immunoprecipitated with protein G and eluted proteins were immunoblotted with anti-CD45 antibody or BTN2A2 antibody. Input is ∼2% of cell lysate. One representative blot is depicted. (D) Ribbon diagram representation of protein-protein interaction between BTN2A2 (pink) and CD45 (PTPRC)(white) showing potential amino acids (blue sticks) determining their interaction. Two residues N419 and N468, key glycosylation sites, located in the fibronectin domain of CD45 that is critical for the interaction is indicated in a spherical model. (E) Western blot demonstrating the expression of CD45 in wild type Jurkat cells (lane 1) and Jurkat cells with CD45 knock-down (CD45-KD) (lane 2); CD45 expression in CD45-KD cells in which wild type CD45 or mutant CD45 (double mutant - N419A and N468A) were expressed is depicted in lanes 3-4. GAPDH was used internal control. (F) Co-immunoprecipitation of CD45 with BTN2A2-Fc or Fc in activated Jurkat cells (1 μg/ml of anti-CD3 antibody for 48hrs) lacking CD45 in which wild type CD45 or mutant CD45 (double mutant - N419A and N468) were expressed as described in Methods and in Panel (E). Following Protein-G pulldown, eluted proteins were immunoblotted with anti-CD45 antibody or BTN2A2 antibody. Input is ∼2% of cell lysate. Panel on the left is one representative experiment and panel on the right is a summary of 3 independent experiments . (G) Co-immunoprecipitation of CD45 with BTN2A2-Fc or Fc in activated Jurkat cells (1 μg/ml of anti-CD3 and anti-CD28 antibody for 48hrs) treated with or without PNGase as described in Methods. Following Protein-G pulldown, eluted proteins were immunoblotted with anti-CD45 antibody or BTN2A2 antibody. Input is 2% of cell lysate. Panel is one representative experiment out of 3 independent experiments. (H) Western blot analysis of CD45 isoforms (CD45RO and RA) following immunoprecipitation with protein G agarose beads of lysates obtained from activated Jurkat cells (1 μg/ml of anti-CD3 and anti-CD28 antibody for 48hrs) in the presence of recombinant BTN2A2-Fc or Fc. Input was used as 2% of total cell lysate. Right panel shows quantification of CD45RO and CD45RA co- immunoprecipitation from three independent experiments. (I) CD45RO expression in resting Jurkat cells and activated Jurkat cells (1 μg/ml of anti-CD3 and anti-CD28 antibody for 48hrs). One of two experiments depicted. (J) Binding of BTN2A2-Fc or Fc alone on activated Jurkat cells (1 μg/ml of anti-CD3 and anti-CD28 antibody for 48hrs). Preincubation of activated Jurkat cells with anti-CD45RO antibody, but not anti-CD45RA antibody, inhibited binding of BTN2A2-Fc on Jurkat cells. One of two experiments depicted. (K) MST analysis of the BTN2A2 binding interaction with CD45RO-Fc. The MST dose response data is obtained by titrating recombinant CD45RO-Fc protein (8 μM to 244 pM) against 20 nM fluorescent labeled recombinant BTN2A2 protein. The MST data analysis show that BTN2A2 interacts with CD45RO-Fc with a binding constant (*K*_D_) of 63.5 nM. The data is represented as mean ± SD of triplicates of one independent experiment and the experiment was repeated three times.

We then performed molecular modeling to identify regions in the extracellular region of CD45 that might interact with BTN2A2. Protein homology modeling showed that BTN2A2 interacted with the extracellular fibronectin domain of CD45 protein and in particular amino acids - Asn-419 and Asn-468 (putative N-glycosylation sites based on their location within N-X-S/T consensus motifs) were critical for this interaction (Fig. 2D). It should be noted that the glycosylation of CD45 at N419 was previously reported by mass spectrometry (Wollscheid et al., 2009); interestingly this glycosylation was detected in activated Jurkat cells, but not in non-activated cells. We then performed site-directed mutagenesis of asparagine to alanine and confirmed that mutant CD45 interaction with BTN2A2 was significantly attenuated (Supplementary Fig. 5A-C and Fig. 2E-F). To confirm that N-glycosylation sites on CD45 were critical for the interaction, we treated Jurkat cells with endoglycosidic enzyme PNGase F (N-Glycosidase F) (Supplementary Fig. 6). As predicted, PNGase treatment significantly attenuated the interaction between CD45 and BTN2A2 in activated Jurkat cells (Fig. 2G).

Based on our observations that BTN2A2 showed enhanced interaction with CD45 in activated T cells, we suspected that BTN2A2 was preferentially binding with the CD45RO isoform, because CD45RO is predominantly expressed in activated T cells (Hermiston et al., 2003) and is upregulated in Jurkat cells after activation (Supplemental Fig. 7). Considering this, we repeated the co-immunoprecipitation assays and immunoblotted the eluted proteins using CD45 isoform specific antibodies (Fig. 2H). These experiments confirmed that BTN2A2 binds CD45RO isoform (the predominant isoform in activated T cells), but did not bind CD45RA. Further, cell surface binding studies confirmed that CD45RO expression goes up during Jurkat cell activation (Fig. 2I). Treatment of activated Jurkat cells with anti-CD45RO antibody, but not anti- CD45RA antibody inhibited cell surface binding of BTN2A2-Fc to activated Jurkat cells (Fig. 2J).

To demonstrate direct binding of BTN2A2 to CD45RO, we performed microscale thermopheresis (MST) using recombinant BTN2A2 and CD45 RO, which demonstrated specific high affinity binding of these two proteins with Kd of 63.5nM (Fig. 2K).

These studies confirm that BTN2A2 specifically and directly binds to CD45RO on activated T cells and posit that activation of CD45RO mediates the functional effects reported for BTN2A2.

### BTN2A2 enhances regulatory T cell expansion and suppresses Th17 cell populations in primary mouse T cells

Activated CD4+ T cells differentiate into several effector cell subsets based on activation events and the cytokine milieu present during activation. Since it was previously reported that BTN2A2 induced Foxp3 expression in CD4+ T cells (Ammann et al., 2013) in short-term culture studies we tested whether T cells activated in the presence of BTN2A2 increased Treg populations in splenocytes from Foxp3-EGFP mice that co-express EGFP when Foxp3 is expressed. We cultured murine T cells (Foxp3+ reporter mice -BALB/cJ strain) in the presence of irradiated, allogeneic mouse splenocytes (CD1 mouse strain) with or without recombinant BTN2A2-Fc. These assays showed a 50% increase in CD4+CD25+Foxp3+ T cells, in BTN2A2-Fc treated cells compared to controls, a similar magnitude to that observed in positive control cultures using exogenous TGF-β (Fig. 3A). Parallel assays showed that recombinant BTN2A2 blocked IL-6/TGF-β/IL-1β-induced production of RORγt, the signature transcription factor of Th17 cells (Fig. 3B). Interestingly, short-term exposure of BTN2A2 to activated CD4^+^ T cells led to less proliferation and enhanced survival, phenotypes noted with a pro-differentiation state and similar to what has been reported with TGF-β (McKarns and Schwartz, 2005) (Fig. 3C-D).

**Figure 3:**
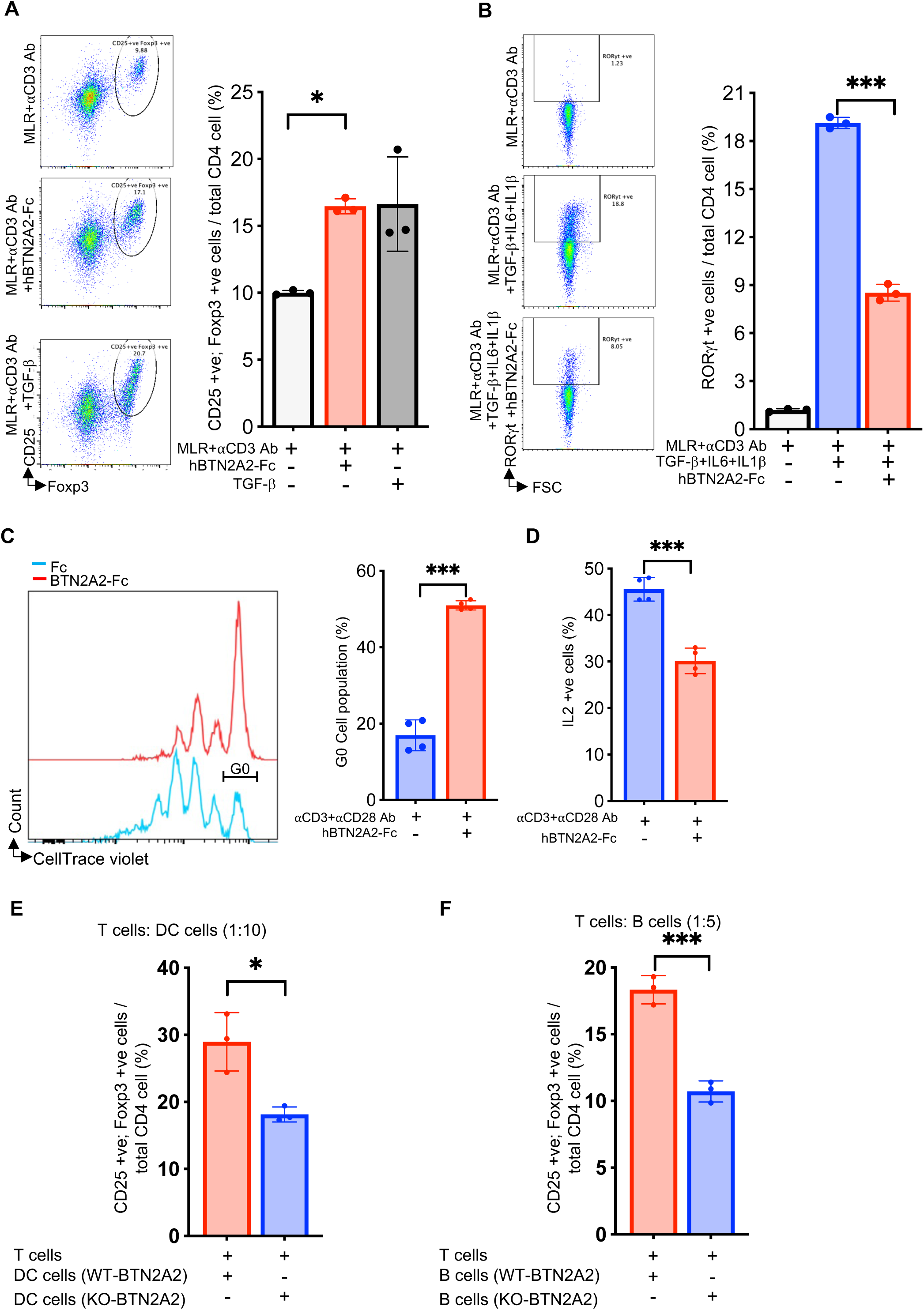
BTN2A2-Fc enhances Tregs and suppress Th17 cell differentiation in invitro mixed lymphocytes reaction (MLR) (A) Flow cytometry analysis plot of primary CD4+ T cells (from spleen and lymph nodes of Foxp3-GFP transgenic mice) incubated with bound 1 μg/ml anti-CD3 and in the presence or absence of 10 μg/ml BTN2A2-Fc fusion protein in MLR for 7 days as described in methods and analyzed for CD4+CD25+ Foxp3-GFP+ve cells expression. As positive control, effects of TGF-β (1 ng/ml) under the same conditions are also depicted. Right panel show summary plot depicted as mean±SD for 3 independent experiments. One-way ANOVA with Tukey’s multiple comparison test; *p*<0.05 (*). (B) Flow cytometry analysis of primary CD4 +T cells (isolated from murine spleen and lymph nodes) incubated for 5 days with anti-CD3 (0.5 μg/ml) antibody alone or anti-CD3 +TGF-β (1.5 ng/ml) +IL6 (10 ng/ml) +IL-1β (10 ng/ml) and/or recombinant BTN2A2-Fc (10 μg/ml) in MLR and analyzed for CD4+RORγt+ve cells. Right panel show summary plot depicted as mean±SD for 3 independent experiments. One-way ANOVA with Tukey’s multiple comparison test; *p*<0.05 (*). (C) Flow cytometry plots of CellTrace violet stained purified mouse CD4+ve cells incubated with immobilized anti-CD3 and anti-CD28 antibody (0.5 μg/ml) in the presence of immobilized recombinant BTN2A2-Fc (10 μg/ml) or Fc (10 μg/ml) for 3 days. Right panels show bar graph depicted as a percentage of non- proliferated cells (G0) (D) Flow cytometry plots of IL2 positive purified mouse CD4+ve cells incubated with immobilized anti-CD3 and anti-CD28 antibody (0.5 μg/ml) in the presence of immobilized recombinant BTN2A2-Fc (10 μg/ml) or Fc (10 μg/ml) for 3 days. Right panels shows bar graph of IL2 positive cells percentage. Data for experiments in (C-D) are depicted as mean ± SD (n=4). Unpaired t-test; p<0.05 (*), p<0.001 (***). (E) Flow cytometry analysis of Foxp3-GFP +ve cell (%) population in total CD4+ T cells co-cultured with dendritic cells (DC) at day 7. Purified T cells were incubated in RPMI ex-vivo for day7 with DC cells (1:10 ratio) isolated from wild- type or BTN2A2-/- mice. (F) Flow cytometry analysis of Foxp3-GFP +ve cell (%) population in total CD4+ cells co-cultured with B cells at day 7. Purified T-cells were incubated ex-vivo for day7 with B-cell (1:5 ratio) isolated from wild-type or BTN2A2-/- mice. Data for experiments in (E-F) are depicted as mean ± SD (n=3). Unpaired t-test; p<0.05 (*), p<0.001 (***).

To further explore the short-term effects of BTN2A2 on T cell subset expression patterns in mixed lymphocyte reaction (MLR) culture conditions, we assessed gene expression patterns for molecules related to Th1, Th2, Treg and Th17 cells by RT-PCR. BTN2A2 upregulated IL2Rbeta and Foxp3 and downregulated IL-21 consistent with our in vitro findings of expansion of Tregs and suppression of Th17 cells (Supplementary Fig. 8A-C). The assays further showed downregulation of transcription factors regulating Th1 (TBX21) and Th2 (GATA3) differentiation pathway at day 5 (Supplementary Fig. 8D-F).

To explore the role of endogenous BTN2A2, we genetically knocked out murine BTN2A2 using CRISPR-Cas9 strategy (Supplementary Fig. 9A-B) and confirmed that expression of BTN2A2 was absent in professional APCs (CD11c+ dendritic cells and CD19+ B cells) in knock-out mice compared to wild-type littermates (Supplementary Fig. 9C-D). Interestingly, when isolated APCs from BTN2A2-/- animals were co-cultured with murine T cells from Foxp3+ reporter mice, absence of BTN2A2 from both isolated dendritic cells and isolated B cells from BTN2A2-/- animals impaired mouse T cell differentiation to Foxp3+ Tregs when compared to dendritic cells and B cells from wildtype littermates (Fig. 3E-F)

### BTN2A2-induced enhancement of Treg/Th17 balance is dependent on CD45 phosphatase activity

To test whether CD45 phosphatase activity was essential for BTN2A2’s actions on TCR signaling and Treg expansion, we used a previously validated CD45-specific phosphatase inhibitor (Perron et al., 2014). Addition of BTN2A2-Fc to activated Jurkat T cells decreased phosphorylation of ZAP70 kinase, while inhibition of CD45 phosphatase activity with this small molecule inhibitor (Perron et al., 2014) abrogated the effect (Fig. 4A). When we inhibited CD45 phosphatase activity during BTN2A2-Fc treatment we also observed complete abrogation of the effects of BTN2A2-Fc on increased Treg and decreased Th17 populations (Fig. 4B-C). Taken together with other experiments described above, these studies link BTN2A2 effects on CD45 phosphatase activity within the TCR complex and immune synapse and Treg and Th17 cellular balance.

**Figure 4:**
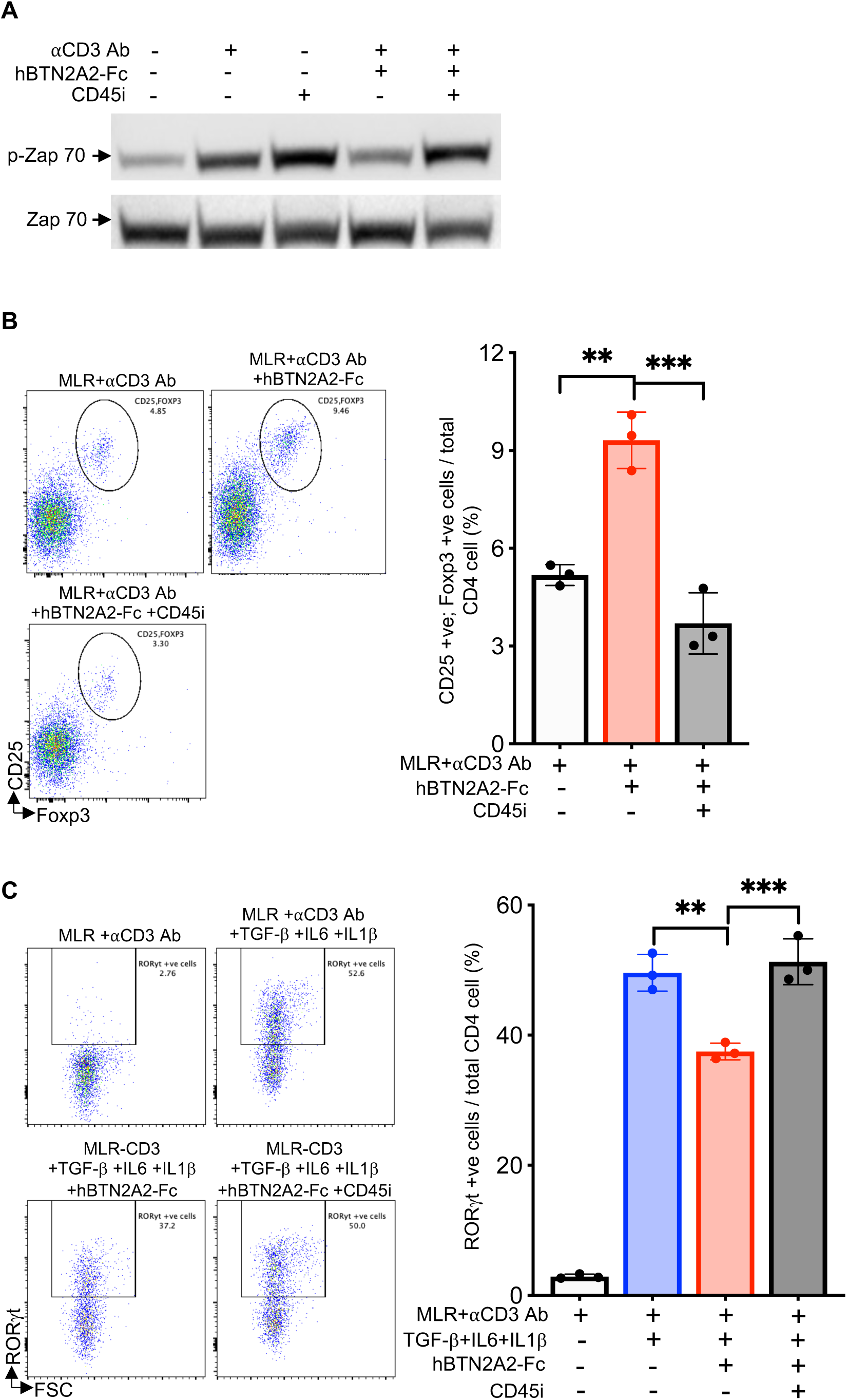
Inhibition of CD45 phosphatase activity in primary immune cells blocks BTN2A2 mediated Treg differentiation and Th17 suppression. (A) Immunoblot analysis of phosphorylated ZAP-70 (p-Zap70) and total Zap70 in Jurkat cells pretreated with CD45 phosphatase inhibitor for 1 hr and stimulated for 3 min with plate-bound anti-CD3 (10 μg/ml) antibody in the presence or absence of recombinant BTN2A2-Fc (10 μg/ml). (B) Flow cytometry analysis plot of primary CD4+ T cells (from spleen and lymph nodes of Foxp3-GFP transgenic mice) incubated with bound 1 μg/ml anti-CD3 and/or 10 μg/ml BTN2A2-Fc fusion protein in MLR for 5 days in absence or presence of CD45 phosphatase inhibitor (125nM) and analyzed for CD4+CD25+ Foxp3-GFP+ve cells expression. Right panel shows summary plot depicted as mean ± SD for 3 independent experiments. (C) Flow cytometry analysis of primary CD4 +T cells (isolated from murine spleen and lymph nodes) incubated for 5 days with anti-CD3 (0.5 μg/ml) antibody alone or anti-CD3 +TGF-β (1.5 ng/ml) +IL6 (10 ng/ml) +IL-1β (10 ng/ml) and/or recombinant BTN2A2-Fc (10 μg/ml) in MLR for 7 days in absence or presence of CD45 phosphatase inhibitor (125nM) and analyzed for CD4+RORγt+ve cells. Right panel shows summary plot depicted as mean ± SD. One-way ANOVA test with Tukey’s multiple comparison test; *p*<0.01 (**), *p*<0.001 (***).

### BTN2A2-Fc therapy exhibits immunoregulatory function in vivo

To test whether and how the in vitro observed immunoregulatory effects of BTN2A2 correlate to findings *in vivo*, we employed two distinct murine immune-mediated model systems. Consistent with previous reports (Saito et al., 2022; Tipping and Holdsworth, 2006), injection of nephrotoxic serum (NTS) in wild type B6 mice induced crescentic glomerulonephritis with significant proteinuria (Fig. 5A-D). Administration of BTN2A2-Fc during the days following NTS administration reduced proteinuria and glomerular crescent formation. Administration of BTN2A2-Fc also increased CD4+ Foxp3 gene expression (Fig. 5E), decreased CD4+ RORγt gene expression (Fig. 5F), attenuated T cell activation marker CD5 levels (Fig. 5G) in CD4+ T cells purified from spleen and lymph nodes. Further, BTN2A2-Fc lowered IL17A protein expression in the kidneys compared to controls (Fig. 5H-I).

**Figure 5:**
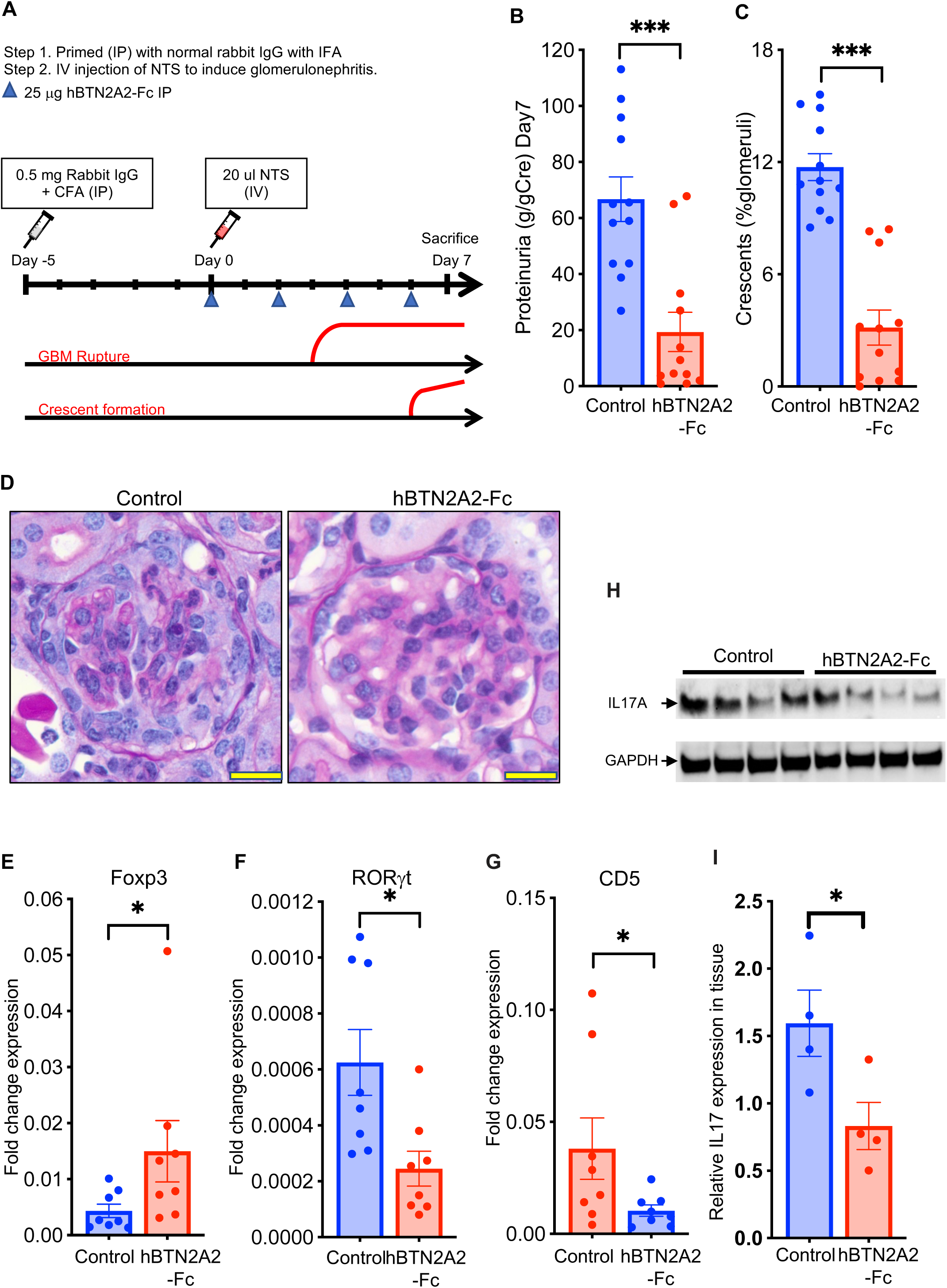
BTN2A2-Fc ameliorates crescentic glomerulonephritis in mice induced by nephrotoxic serum (NTS) (A) Schematic protocol of NTS-induced nephrotoxic glomerulonephritis model and histological changes. GBM = glomerular basement membrane. (B) Quantification of mean ± SEM proteinuria (on Day 7) in mice with NTS induced glomerulonephritis treated with recombinant BTN2A2-Fc protein or vehicle control (control) (N=12/group); Mann-Whitney p<0.001(***) (C-D) Representative images of glomerular injury by PAS staining. Crescent formation in mice with NTS induced glomerulonephritis treated with recombinant BTN2A2-Fc protein or control (D). Scale bars are 25 μm. Summary panel (C) depicts quantification of % glomeruli with crescents (mean ± SEM, N=12/group); Mann-Whitney p<0.001(***) (E-F) Relative Foxp3 (E) and RORγt (F) expression compared to GAPDH as internal control in CD4+ T cells purified from spleen and lymph nodes from mice with NTS induced glomerulonephritis treated with BTN2A2-Fc protein or control. Data depicted as mean ± SEM (N=8/group). Mann-Whitney test; *p*<0.05 (*). (G) Quantitative PCR analysis show CD5 expression in CD4+T cells from mice with NTS induced glomerulonephritis treated with BTN2A2-Fc protein or control.. N=8/group (H-I) Immunoblot analysis of IL17A in kidney tissue lysate of mice with glomerulonephritis treated with recombinant BTN2A2-Fc or control (H). Panel I shows quantification of western blot. Data represented as mean ± SEM. N=4 per group. Unpaired t-test; *p*<0.05 (*).

We next administered a lower dose of NTS (50% less than the amount used in experiments above) to BTN2A2-/- mice (Supplementary Fig. 9) and wildtype littermate controls (Fig. 6). These experiments demonstrated that lower doses of NTS induced relatively mild crescentic glomerulonephritis and proteinuria among wild-type control animals, however, in BTN2A2-/- animals we saw an exacerbation of crescentic glomerulonephritis and severe proteinuria (Fig. 6A-C). We also observed reduced Foxp3 and enhanced RORγt gene expression in splenic/lymph node CD4+ T cells from the BTN2A2-/- mice vs. controls (Fig. 6D-E). CD5 was enhanced in CD4+ T cells in BTN2A2-/- mice compared to wild type mice (Fig. 6F). Analysis of kidney tissue one week after administration of nephrotoxic serum also showed increased IL17A protein expression in the kidneys of BTN2A2-/- vs controls animals (Fig. 6G-H).

**Figure 6:**
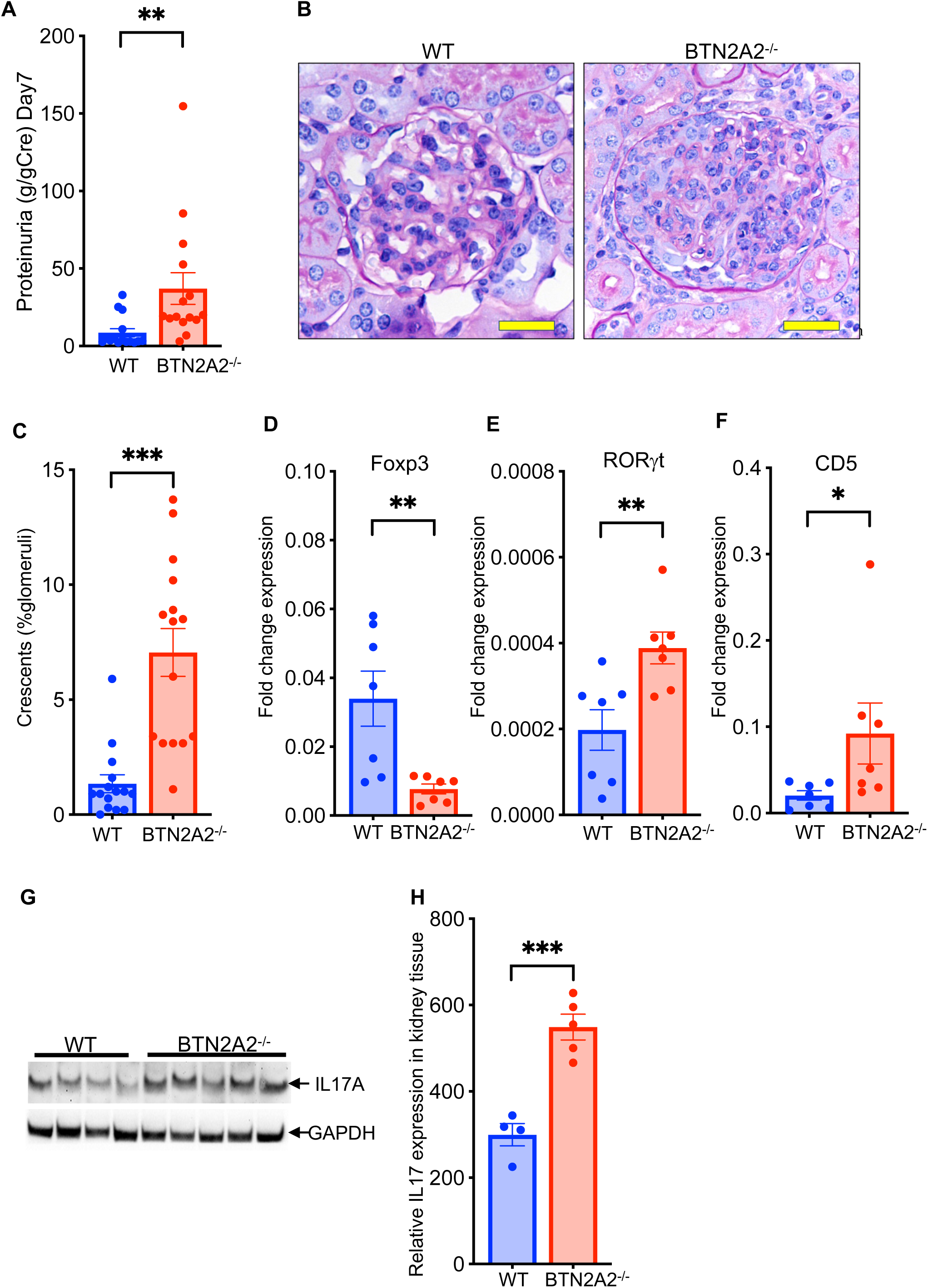
BTN2A2 (-/-) mice show exacerbated crescentic glomerulonephritis. (A) Scatter plot shows quantification of mean ± SEM of proteinuria in wild-type and BTN2A2 (-/-) mice with nephrotoxic glomerulonephritis on Day 7 (N = 15). Mann-Whitney test; *p*<0.01 (**). (B, C) Representative images of glomerular injury by PAS staining. Wild-type mice show minimal glomerular injury with a low dose of nephrotoxic serum, while BTN2A2 (-/-) mice show severe glomerulonephritis (B). Scale bars are 25 μm. Summary data are depicted as mean ± SEM of %Glomerular crescents (C). N=15 per group for all experiments. Mann-Whitney test; *p*<0.001 (***). (D-E) Relative Foxp3 mRNA (D) and RORγt mRNA (E) expression compared to GAPDH as internal control in CD4+ve T cells purified from spleen and lymph nodes cells from wild-type and BTN2A2-knockout mice with nephrotoxic serum induced glomerulonephritis. (F) Quantitative PCR analysis show CD5 expression in CD4+T cells from wild-type and BTN2A2-knockout mice with nephrotoxic serum induced glomerulonephritis. N=7/group (G-H) Immunoblotting for IL17A expression in kidney tissue lysate using anti-IL-17A antibody (G) and quantitation (H) from wild-type and BTN2A2 null mice with nephrotoxic serum induced glomerulonephritis. n≥4 per group. Unpaired t-test; *p*<0.01 (**), *p*<0.001 (***).

To evaluate the generalizability of the role of BTN2A2 in immunoregulation, we studied the effects of recombinant BTN2A2-Fc in DBA/2 mice mated with CBA/J mice, a model of immunologically mediated abortions (Clark et al., 2008). We confirmed that DBA/2 male x CBA/J female had reduced litter sizes and higher spontaneous adsorption rates compared to CBA/J male x DBA/2 females. Administration of BTN2A2-Fc improved litter size and rescued the excess abortion rates that were noted in the DBA/2 male x CBA/J female (Fig. 7A-C). The beneficial effects of BTN2A2-Fc were associated with increased frequencies of splenic/lymph node Foxp3+ Tregs, reduced frequencies of splenic/lymph nodeTh17 cells and attenuation of CD5, consistent with our findings in the autoimmune GN model (Fig. 7D-G). In addition, BTN2A2-Fc protein also reduced placental IL-17 protein expression that correlated with improved litter sizes (Fig. 7H-I).

**Figure 7:**
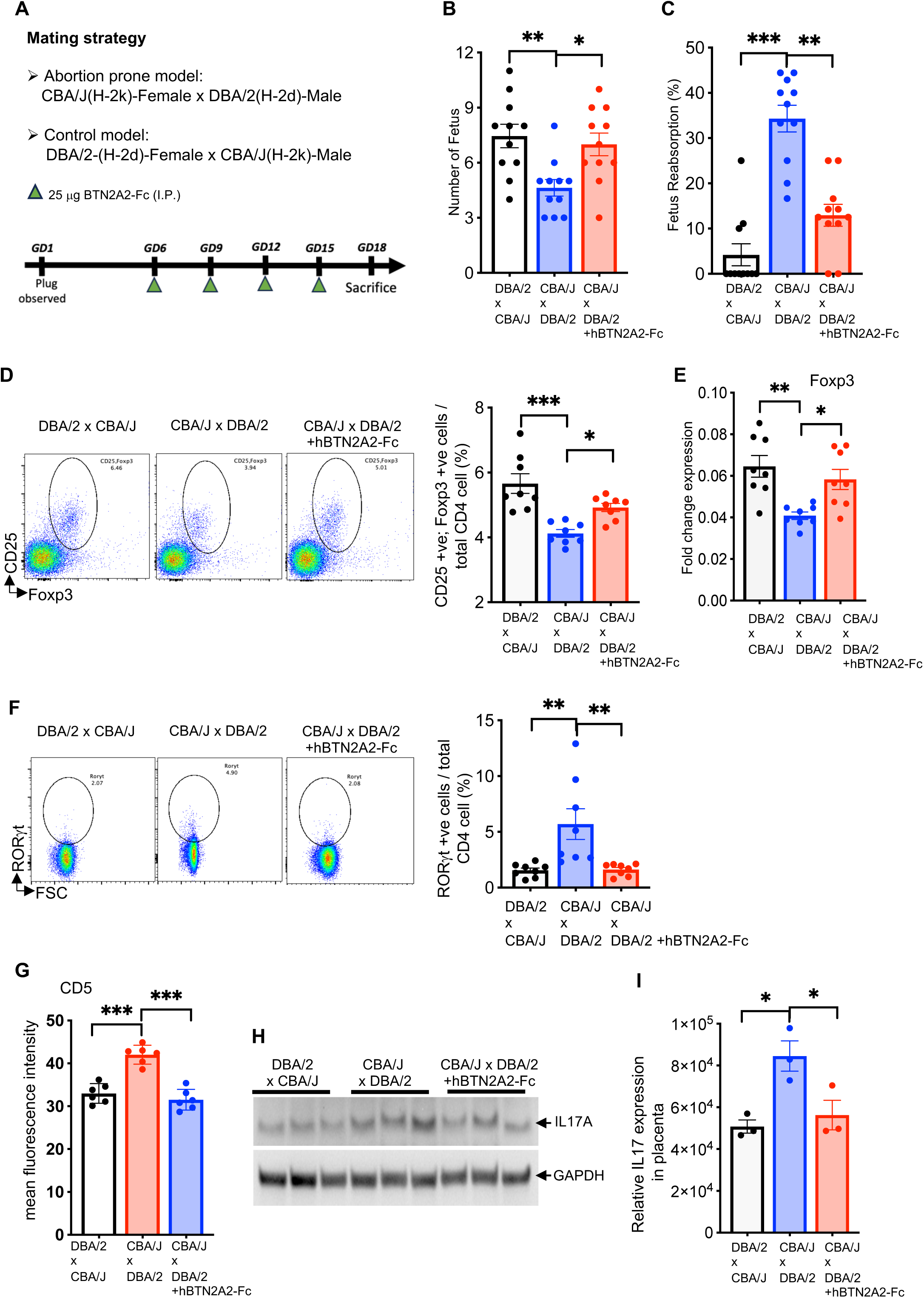
BTN2A2-Fc rescues fetus resorption and increases litter size in CBA/J x DBA/2 pregnancy model. (A) Schematic of the Abortion prone model. (B-C). Pregnant CBA/J x DBA/2 mice treated with BTN2A2 throughout pregnancy has improved litter size (B), reduced resorption (C) compared to untreated mice. As additional control, DBA/2 x CBA/J mice has been included. Data (B-C) represented as mean ± SEM. N=11 per group for all experiments. Kruskal-Wallis test with Dunn’s test for multiple comparison; *p*<0.05 (*), *p*<0.01 (**), *p*<0.001 (***). (D-E) Flow cytometry (D) Quantitative PCR (E) analysis show increase in Foxp3 expressing CD4+T cells in pregnant CBA/J x DBA/2J mice treated with recombinant BTN2A2-Fc as compared to control mice. N=8/group (F) Flow cytometry analysis (left panel) show reduced number of CD4+RORγt +ve cells in pregnant CBA/J x DBA/2J mice treated with recombinant BTN2A2-Fc as compared to control mice. Right panel shows quantification of the summary data. N=8/group (G) Flow cytometry analysis show mean fluorescence intensity of CD4+CD5 +ve cells in pregnant CBA/J x DBA/2J mice treated with recombinant BTN2A2-Fc as compared to control mice DBA/2J x CBA/J. N=6/group (H, I) Immunoblot analysis of IL17A in placental tissue lysate of pregnant CBA/J x DBA/2J mice treated with recombinant BTN2A2-Fc or control (left panel). Right panel shows quantification of summary data. N=3 per group One-way ANOVA test with Tukey’s multiple comparison test for experiments in C-G; *p*<0.05 (*), *p*<0.01 (**), *p*<0.001 (***).

### BTN2A2 enhances Treg cell expansion and suppresses Th17 cell populations in human PBMCs

To evaluate whether the effects of recombinant BTN2A2 apply to human T cells, we analyzed human PBMCs activated by mixed lymphocyte reactions (MLR, using allogeneic stimulator cells) with and without recombinant BTN2A2-Fc, and quantified CD4+CD25+CD127^low/-^ Foxp3+ Treg cell numbers 7 days later. These co-cultures showed that the BTN2A2-Fc induced 2-fold expansion of Tregs under MLR conditions compared with controls; however BTN2A2-Fc was unable to induce Tregs in the presence of CD45 inhibitor (Fig. 8A-B). Parallel experiments revealed that recombinant BTN2A2-Fc robustly blocked the TGFβ-IL6-IL1β induced increase in Th17 cells (Fig. 8C-D). Interestingly, BTN2A2-Fc was unable to block cytokines induced Th17 cells in the presence of CD45 inhibitor. Taken together, these observations suggest that CD45 phosphatase activity is necessary for BTN2A2’s actions in humans as well.

**Figure 8:**
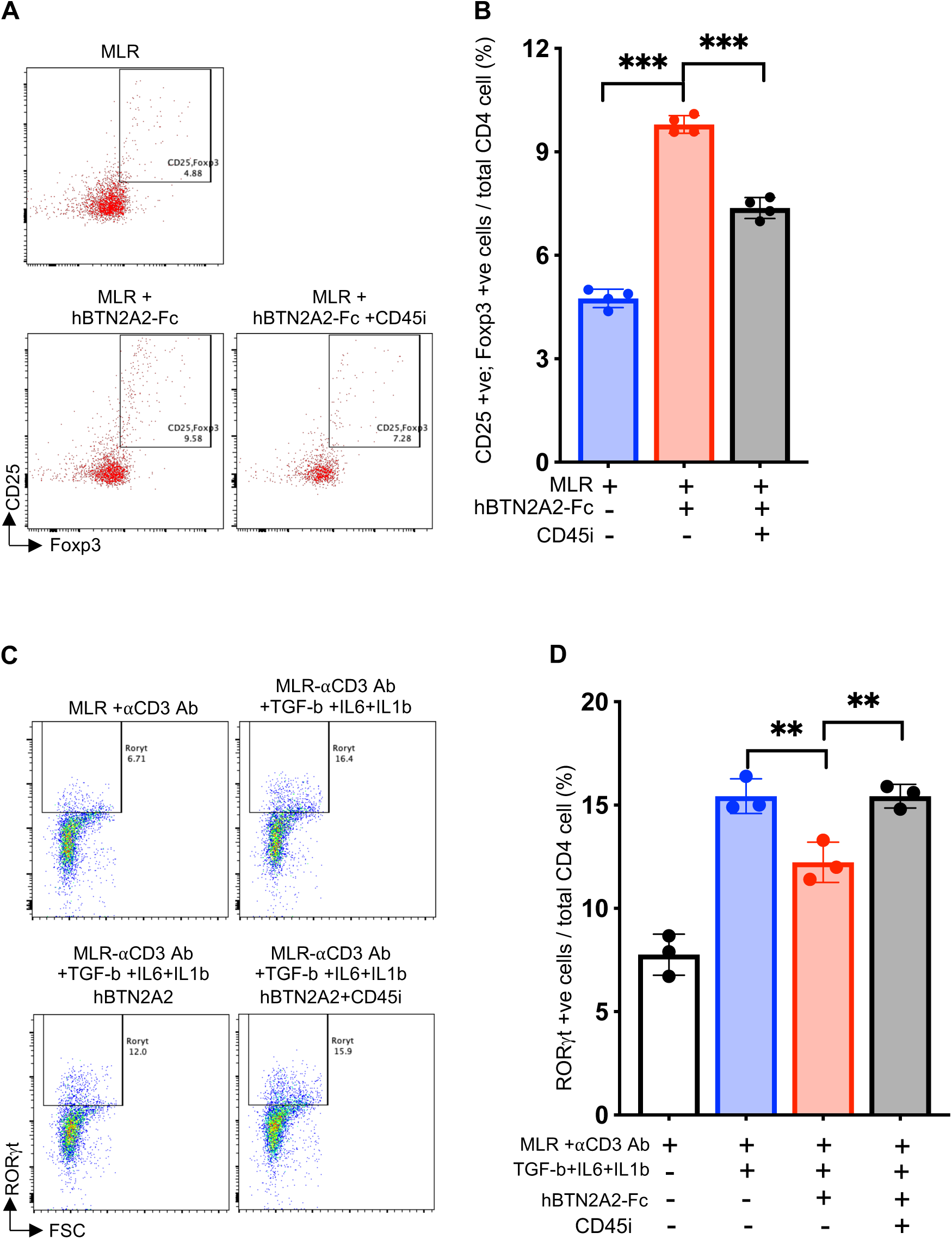
BTN2A2-Fc enhances Tregs and suppress Th17 cells differentiation in in vitro human PBMC mixed lymphocyte reactions. (A-B). Flow cytometry plots showing percentage of Treg cells (CD4+CD25+Foxp3 +ve cells) in human PBMC incubated with /without recombinant BTN2A2-Fc (10 μg/ml) in MLR for 7 days in absence or presence of CD45 phosphatase inhibitor (125nM). Data depicted as mean +/-SD showing percentage of Treg cells (CD4+CD25+Foxp3 +ve cells) among total CD4+ cells. N=4 per group for all experiments. Unpaired 2-tailed *t* test; *p*<0.001 (***). (C-D). Flow cytometry plots showing percentage of CD4+RORγt +ve (Th17 cells) in human PBMC incubated with anti-CD3 antibody (0.5 μg/ml) only or anti-CD3 antibody +TGF-β (1.5 ng/ml) +IL-6 (10 ng/ml) +IL-1β (10 ng/ml) in absence or presence of recombinant BTN2A2-Fc (10 μg/ml) in MLR for 5 days in absence or presence of CD45 phosphatase inhibitor (125nM). Data represent mean ± standard deviation. N=3 per group for all experiments. One-way ANOVA test with Tukey’s multiple comparison test; *p*<0.01 (**).

## DISCUSSION

Data presented here provide multiple lines of evidence supporting the conclusion that BTN2A2 functions as a ligand for CD45, preferentially binding to CD45RO isoform, and that it functions to prevent segregation of CD45 from the TCR/CD3 complex, resulting in sustained CD45’s phosphatase activity within the immune synapse which reduces downstream signaling cascade (Fig. 9). Consequently, BTN2A2 reduces proliferation of effector Th17 cells while promoting differentiation of T regulatory cells, and reduces severity of NTS-induced glomerulonephritis and immune mediated pregnancy loss in mouse models of immunologic diseases. Together with previous work showing that

**Figure 9:**
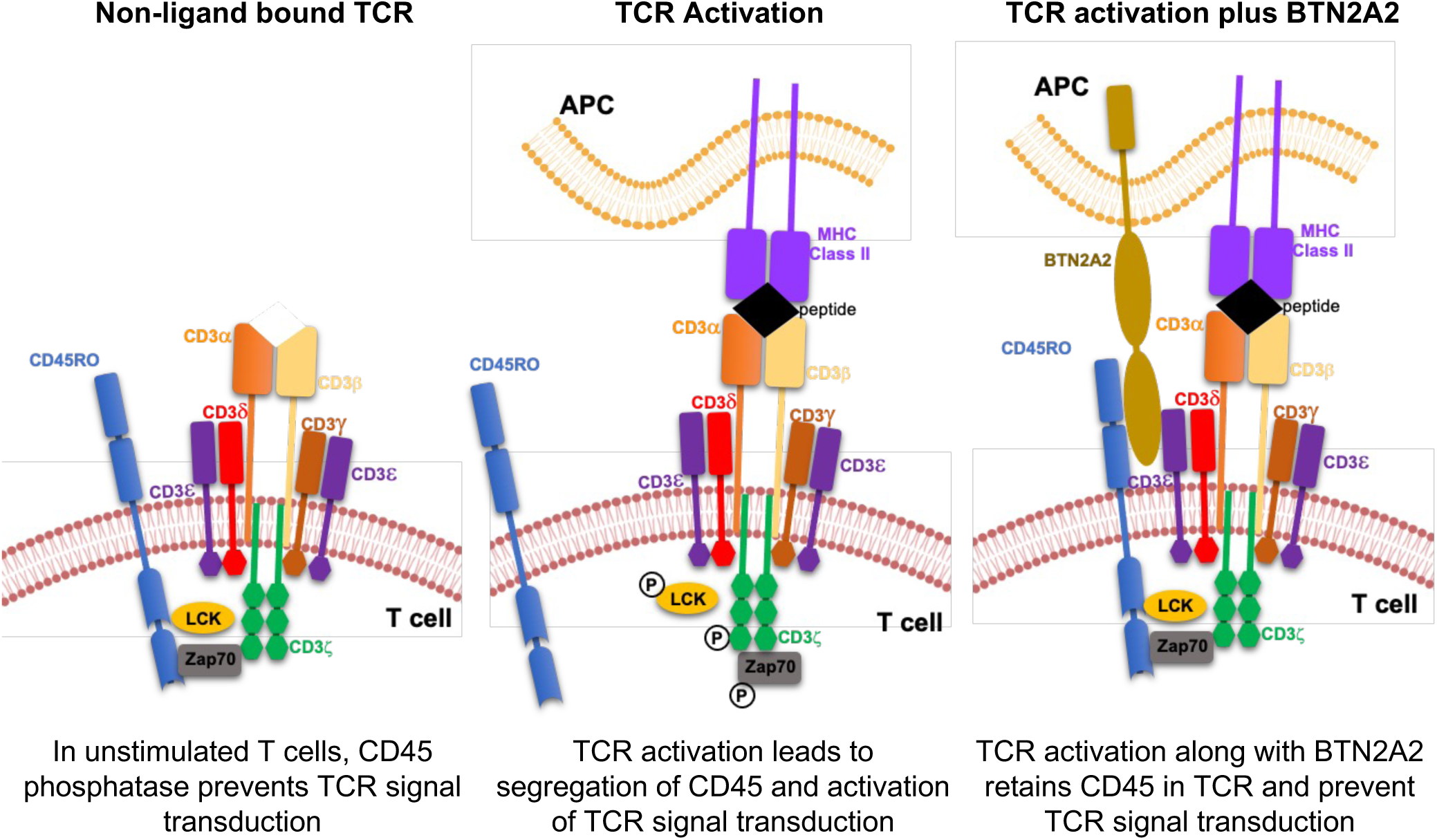
Schematic of the model by which BTN2A2 as a co-inhibitor of TCR signaling via CD45 phosphatase.

APC-derived BTN2A2 regulates T cells (Sarter et al., 2016), our findings suggest that during cognate T cell/APC interactions, APC-presented BTN2A2 ligates T cell-expressed CD45RO to the TCR and functions as a molecular brake at the time of activation by disrupting CD45 segregation from the TCR complex (an event previously implicated as necessary for T cell activation) (Leupin et al., 2000). Our data are also consistent with Payne et al who recently reported that BTN3A1, another family member of BTN proteins, inhibits tumor reactive T cell response by preventing segregation of N-glycosylated CD45 from the immune synapse (Payne et al., 2020). We confirm that BTN2A2 inhibits T cell activation by interacting with CD45RO, that specific N-glycosylated residues in CD45 (Asp-419 and Asn-468) are critical for this interaction and confirm that point mutations of these residues or PNGase treatment significantly attenuated the CD45/BTN2A2 interaction. Our data suggest that in activated T cells, BTN2A2 binds to both CD45 and CD3e, likely maintaining them in proximity, enabling CD45 phosphatase activity to persist, enabling dephosphorylation of CD3 ITAM domains and Zap70 and limiting downstream TCR signals required for full T cell activation (Fig. 9).

Our data indicate that BTN2A2 enhances CD45’s phosphatase activity within the TCR complex which modulates the TCR signaling but does not fully prevent it, and consequently, permits expansion of Tregs while reducing Teffector cell differentiation/expansion. In prior studies, transient targeting of CD45 induced potent antigen-specific regulatory T cells and induced transplant tolerance (Camirand et al., 2014; Picarda et al., 2017). Our data suggests that BTN2A2 expressed on APCs and DCs is likely the endogenous ligand for CD45 on T cells. CD45 is expressed as multiple spliced products and it is currently believed that CD45RA and CD45RB are replaced by CD45RO following T cell activation (Hermiston et al., 2003). Our structural and functional studies suggest that BTN2A2 predominantly binds to the CD45RO isoform that is expressed on activated T cells, signifying specificity for BTN2A2’s actions (Hermiston et al., 2003). BTN2A2 bound directly to CD45RO with kD of 63.5nM which is in the range of binding affinities proposed for other B7 family members with receptors CD28 and CTLA-4 (Collins et al., 2002). Our findings also support prior work where targeting naïve animals with anti-CD45RB antibody resulted in increased expression of CD45RO isoforms that correlated with upregulation of immunoregulatory T cell subset (Basadonna et al., 1998). While multiple human glycoproteins, including galectin-1 and CD22, were shown by others to bind CD45 (Alborzian Deh Sheikh et al., 2021; Earl et al., 2010), there is no clear evidence that these putative ligands modulate CD45 phosphatase activity. Other identified ligands for CD45 include pUL11, a viral protein expressed by cytomegalovirus-infected cells, as well as E3/49K protein expressed by adenovirus-infected cells; both of these viral proteins may play roles in inhibiting anti- viral immunity by inhibiting TCR signaling, although precise mechanisms remain unclear (Windheim et al., 2023; Zischke et al., 2017).

We showed that BTN2A2-Fc limits clinical manifestations of glomerular crescent formation and proteinuria in murine NTS glomerulonephritis and reduces miscarriages in a distinct orthogonal mouse model, both of which were associated with augmented Treg and reduced Th17 cell populations. Consistent with prior data CD5, a T cell activation marker was attenuated by BTN2A2-Fc (Frech et al., 2023). Together with previous work in murine experimental autoimmune encephalomyeletis (Sarter et al., 2016), our data further indicate that BTN2A2-Fc is a potentially useful immune modulator with translational potential in a variety of immune-mediated diseases and transplantation. Our observations are an important step forward in understanding how naturally occurring proteins can modify inflammatory events by interacting with CD45 to sustain TCR complex phosphatase activity, increasing Treg and decreasing Th17 cell populations.

Our work leaves unanswered questions and paths for future work. How does BTN2A2 regulate autoimmunity? While we show that TCR signaling through ZAP70 is modulated by BTN2A2 in the presence of aCD3, we do not know how this dampening of TCR signaling leads to up-regulation of Tregs and down-regulation of Th17 signaling.

Prior studies from other groups have also suggested that switching the antigen stimulation from an acute high level to chronic low-level stimulation may lead to more Treg cells and immune tolerance (Gottschalk et al., 2010; Kretschmer et al., 2005).

CD45 has been proposed as a signaling gatekeeper in T cells (Courtney et al., 2019), however structural and functional studies are now needed to precisely define how enabling graded signaling outputs in the presence or absence of BTN2A2 leads to promotion tolerogenic signals. Further understanding the changes in the CD45 isoform and glycosylation status following activation may offer new immune regulatory options (Earl and Baum, 2008). We also do not know whether this pathway of BTN2A2 mediated Treg upregulation is dependent on TGF-β signaling. In experimental studies, CD28-B7 blockade with CTLA4-Ig can prevent the development of autoimmune glomerulonephritis (Reynolds et al., 2000). It would be interesting to evaluate whether BTN2A2 can synergize with co-stimulatory pathway molecules such as CTLA4-Ig that also promote tolerance by upregulating T regs (Guillot et al., 2000). Other butyrophilin family members such as BTN2A1 and BTN3A1 are known to be important for phosphoantigen mediated γδ T cell activation (Rigau et al., 2020). Additional studies will be needed to determine whether BTN2A2 may also regulate γδ T cell activation.

Furthermore, since BTN2A2 is the only butyrophilin family member expressed by professional APCs, including antigen-specific B cells, further studies are needed to investigate whether BTN2A2 plays a role in communicating antigen-specific immune tolerance between autoantigen-specific APCs and CD45RO-expressing autoantigen-specific cognate memory T cells. Since CD45RO is also expressed on B cells, it is possible that some of the beneficial effects of BTN2A2 in our animal studies may be derived from its actions on B cells. In this regard, Szodoray et al. demonstrate that T helper signals upregulate CD45 phosphatase activity in B cells. High CD45 phosphatase activity in memory B cells controls their effective differentiation toward antibody-secreting cells in response to T helper signals (Szodoray et al., 2021).

Additional work is needed to determine whether the immunomodulatory properties of BTN2A2 may act in cis (Hui, 2023), for example in memory B cells that express high levels of ligand and receptor.

In conclusion, our findings suggest that BTN2A2 ligation of CD45 phosphatase to the TCR complex leads to dampened TCR signaling resulting in expansion of Tregs and suppression of Th17 cells. While we demonstrate a beneficial effect of BTN2A2 in two orthogonal models of autoimmunity/immune tolerance, our data suggest that BTN2A2/CD45 signaling pathway could be targeted in a wide variety of immune-mediated diseases such as inflammatory bowel disease, rheumatoid arthritis, multiple sclerosis, and transplant rejections. A better understanding of the precise targets of BTN2A2 based on its modes of action is critical for the development of new treatment strategies in autoimmune and chronic inflammatory diseases and transplant rejection.

## MATERIALS AND METHODS

### BTN2A2-Fc construct generation and purification of recombinant protein

Soluble recombinant BTN2A2-Fc protein was generated using baculovirus infected insect cell system, a method that is both scalable and has been successfully used for generation of recombinant proteins in the immune system (Hilton et al., 2015). An upstream 711 bp region of human BTN2A2 gene (NP_008926.2) containing signal peptide and two extracellular domains (IgV and IgC2) was amplified using specific primers (See supplementary Table 1) from cDNA from 293T cells. Amplified region was first cloned into pFUSE-hIgG1-Fc1 vector (InvivoGen Catalog # pfuse-hg1fc1) using Age1 and EcoRV restriction sites. Subsequently, BTN2A2 region along with Fc region was amplified using specific primers (Supplementary Table 1) and cloned into pBacPak8 vector using Xba1 and Sac1 restriction sites. Paired-end sequencing of the clone was performed by sanger sequencing to confirm the sequence. Primers were synthesized from Integrated DNA Technologies, Inc (IDT) and Phusion® High-Fidelity PCR Master Mix with HF Buffer was used in all PCR amplification reaction (New England Biolabs). Baculovirus were generated by co-transfecting pBacPak8-BTN2A2- Fc clone with linearized baculovirus DNA (TakaraBio Catalog# 631401) into Sf9 insect cells according to manufacturer protocol. Preparation of baculovirus passage (P0, P1, P2) was done as described in manufactures protocol (TakaraBio, Cat # 631402). P3 baculovirus stock was used for large scale protein purification using Hi5 cells grown in serum free media (Express Five™ SFM, ThermoFisher, Cat# 10486025). Pierce™ Protein G Agarose columns (ThermoFisher Scientific, Cat# 20398) was used for purification of secreted soluble-BTN2A2-Fc protein from the supernatant. Purified protein was finally suspended in 1x phosphate buffer saline (1xPBS). Protein purity was checked by SDS-PAGE followed by Coomassie blue staining. Specificity of the purified protein was confirmed by western blot analysis using specific BTN2A2 antibody.

Commercially available recombinant 293 cells derived human BTN2A2-Fc (Cat# 8918- BT; R&D systems) and mouse BTN2A2 (Cat# 8997-BT-050; R&D systems) was used for comparison studies with baculoviral-generated recombinant B2NT2A2-Fc. For some biochemical studies, we also generated BTN2A2-His tag protein (without Fc tag) in Hi5 cells as described above and purified using Ni-NTA Purification System (Invitrogen cat# K95001).

### T-cells activation and IL-2 secretion assay

In vitro T cell activation was performed in Jurkat cells as described previously (Nguyen et al., 2006). Briefly, 96-well plates were coated with 1μg/ml concentrations of anti-CD3 mAb (clone OKT3) in PBS at 4°C overnight in absence or presence of recombinant BTN2A2-Fc (10 μg/ml) or Fc tag protein (Cat#10702-HNAH, Sino Biological Inc) or 293 cells derived human BTN2A2-Fc or mouse BTN2A2. A total of 2.5x10^5^ Jurkat cells/well were added to precoated flat-bottom 96-well plates. Cells were incubated for 24 hrs in 37°C incubator with 5% CO_2_. Total 100 µl of medial was used to measure IL-2 production using an IL-2 ELISA kit from Millipore-Sigma (Cat# RAB0286-1KT) according to the manufacturer’ protocol.

### In vitro TCR stimulation and signal transduction analysis

For short-term activation of the Jurkat cells (Arnett et al., 2007), 48-well tissue culture plate was coated with anti-CD3 mAb (OKT3, 10μg/mL) and BTN2A2-Fc (10μg/mL) or Fc-Tag (10μg/mL). Plate was incubated overnight at 4⁰C and each well was washed twice with PBS. A total of 2x10^6^ cells in 75μl 1xPBS were then added to each well for 3 min at 37⁰C. The reaction was then stopped with ice cold PBS. Pervanadate was prepared by incubating vanadate (200mM) and H_2_O_2_ (200mM) in 1:2 ratio for 15 min at room temperature. Jurkat cells in 1xPBS were treated with pervanadate at final concentration of 0.1mM for 5 min in incubator (5% CO2 and 37°C). The cells were then collected, lysed in lysis buffer (RIPA or IP lysis buffer) and subsequently subjected to immunoprecipitation reaction or immunoblotting.

### Immunoprecipitation and Immunoblotting

Jurkat cells were activated with plate-bound anti-CD3/anti-CD28 with BTN2A2-Fc or Fc-tag protein or studied un-activated. Cells were collected then washed in PBS and resuspended in IP lysis buffer (10mM HEPES pH 7.5, 0.5mM EDTA, 0.5% NP-40, 250mM NaCl, 1x phosSTOP and protease inhibitors), incubated on ice for 30 min with vortexing, and cleared by centrifugation at 15,000g for 10 min. An aliquot of protein lysate was used for western blotting, while the remainder were incubated for 24 hours at 4⁰C for 8-12 hours with specific antibodies or IgG as described in figure legends.

Protein A/G Agarose (ThermoFisher Scientific) was added to lysate and incubated for additional 2 hrs. Beads were washed once in IP-lysis buffer with 0.5% NP-40 and twice in IP-lysis buffer with 0.2% NP-40. Protein lysate prepared after Jurkat cell treatment and/or immunoprecipitation was separated in 4-12% Bis-Tris gel (ThermoFisher Scientific) and transferred to nitrocellulose membrane. Membrane was blocked for 1 h in blocking solution (5% BSA in PBS with 0.1% Tween [PBST]). The membrane was incubated with primary antibodies overnight at 4°C, washed with PBST three times (10 minutes each), incubated with secondary antibodies for 1 hour at room temperature, and finally washed with PBST three times (10 minutes each). The blots were visualized using an ECL assay (ThermoFisher Scientific). Western blots were probed with anti-phospho-Zap70 anti-phospho- CD3ζ antibody and also antibody against total Zap70 and CD3ζ protein (See Supplementary Table 2 in Supplementary Materials).

For phosphatase activity, immunoprecipitated material was directly mixed in Fluorescein Diphosphate, Tetraammonium Salt substrate (FDP, Catalog # F2999, Thermofisher Scientific) and activity was measured using Molecular Devices SpectraMax® M2 plate readers as recommended by protocol included by manufacturer.

For long-term activation studes, Jurkat cells in suspension were activated with anti- CD3/anti-CD28 antibody (1 μg/ml each) for 48-72 hrs in RPMI medium supplemented with 10% FBS. Jurkat cells were collected, washed with PBS and incubated on ice with 10 μg/ml BTNT2-Fc or Fc protein for 1-2 hours prior to immunoprecipitation studies.

Extracellular crosslinking with BS3 (bis[sulfosuccinimidyl] suberate) was performed according to user manual (Thermo Scientific MAN0011240). BS3 (ThermoFisher) was added at a final concentration of 5 mM and incubated for 30 min at room temperature. The reaction was stopped using Tris HCl, pH 7.5, at a final concentration of 20 mM for 15 minutes at room temperature. The cells were then washed extensively in PBS and resuspended in IP-lysis buffer and cleared by centrifugation at 14,000g for 10 min. Protein G Agarose beads were used to pulldown BTN2A2-Fc and Fc-tagged protein. Beads were washed and eluted with SDS sample buffer supplemented with dithiothreitol. Eluted samples were analyzed by western blot probed with anti-CD45 antibody or BTN2A2 antibody (See Supplementary Table 2).

### Molecular modeling of BTN2A2 and PTPRC (CD45)

A molecular model of BTN2A2 was generated using homology modeling. A search for the homologous structure of the extracellular domain of BTN2A2 revealed that BTN3A2 shares about 47% of sequence homology with BTN2A2. Using the crystal structure of BTN3A2 as a template, the three-dimensional model of BTN2A2 was generated using SWISS-MODEL workspace (Arnold et al., 2006). Subsequently, the molecular structure of BTN2A2 was subjected to a short 1.2ns molecular dynamics simulation using Desmond (Schrodinger, Inc., San Diego, CA). The three-dimensional structure of PTPRC was retrieved from the protein data bank (PDB code: 5FN7) (Chang et al., 2016). To determine the interaction between BTN2A2 and PTPRC proteins, RosettaDock 4.0 was used (Lyskov et al., 2013; Marze et al., 2018). The top 10 predicted models were then subjected to 1.2ns molecular simulation followed by minimization using Desmond. Then, the most energetically stable docking model was used to identify potential amino acids for mutation.

### Microscale Thermophoresis measurement of the BTN2A2 binding interaction with CD45RO-Fc

The binding interaction between BTN2A2-His and CD45RO-Fc was measured by Microscale Thermophoresis (MST) using the Monolith NT.115 instrument (Nanotemper Technologies, München, Germany) as described (Wienken et al., 2010). Briefly, the BTN2A2-His protein was fluorescently labeled by covalent labeling method using the Monolith series protein labeling kit RED-NHS 2nd generation (Amine Reactive; Cat# MO-L011). The 1x PBS buffer supplemented with 0.05% Tween-20 was used as the MST assay buffer to perform the binding experiments. The non-labeled CD45RO-Fc protein (Catolog # 10642-CD, R&D systems, MN) was titrated in the concentration range of 8 μM to 244 pM against the 20nM of fluorescent labeled BTN2A2. The MST data were collected at settings including medium (40%) MST power, 20% LED/excitation power, Nano-RED excitation type and 25°C thermostat setpoint. The BTN2A2-His binding affinity to CD45RO-Fc was determined by fitting the MST response data to the Kd model in the MO Affinity Analysis Software version v2.3 (Nanotemper Technologies, München, Germany).

### CRISPR-Cas9 deletion of CD45 gene and site directed mutagenesis of CD45

CD45 knockdown cell lines were generated using the CRISPR-Cas9 system. Two different guide RNAs were designed complementary to upstream of CD45 exon-1 and downstream to exon-3 based on its protospacer adjacent motif (PAM) sequence (Supplementary Table 1). These guide RNA sequences were cloned into lenti-Crispr v2 blast vector containing the Cas9 coding gene and a blasticidine resistant gene. Lentivirus were generated using envelop and packaging plasmid. Jurkat cells were then transduced with lentivirus expressing guideRNA and Cas9 gene and selected for blasticidine resistant cells. Single-cell cloning was performed to isolate independent clones. Deletion of Exon- 1 to Exon-3 was confirmed by direct Sanger sequencing (Supplementary Figure 3A-B).

CD45 (PTPRC) (NM_002838) Human Tagged ORF Clone was purchased from origene technologies. Inc. Site-directed mutagenesis were performed using the QuickChange mutagenesis kit (Invitrogen), to introduce the N419A and N468A mutations. Introduction of the mutation was confirmed by direct sequencing (Supplementary Figure 3C). Lentivirus were generated using envelop and packaging plasmid. Jurkat cells deleted for CD45 gene were then transduced with lentivirus expressing mutant CD45 gene. For PNGase treatment, Jurkat cells were activated with anti-CD3/anti-CD28 antibody 1μg/ml for at least 48 hours in suspension. Cells were collected and resuspended in 1x glycobuffer-2 along with PNGaseF (100U/10^6^ cells, NEB Cat# P0709S) for 4-8 hrs at 37°C in incubator. Cells were lysed in IP lysis buffer for co-IP assay or used directly for Lectin binding assay using flow cytometry analysis (Supplementary Figure 3D). Ulex europaeus (gorse, furze, Sigma-Aldrich Cat#L5505) was purchased, conjugated with PE dye and used 0.7μg/ml for staining.

### In vitro Treg and Th17 differentiation by mixed lymphocyte reaction (MLR)

For Treg differentiation, fresh spleen, and lymph nodes (inguinal, brachial and axillary) were harvested from mice and washed with RPMI media. Spleen was excised into small pieces and single cells were prepared by mashing the tissue with the plunger end of syringe through a 70 micron strainer and cells were suspended in 5 ml of RPMI media. Cells were stimulated with plate bound anti-CD3 (1μg/ml) with or without BTN2A2-Fc fusion protein (10μg/ml) and co-cultured with CD1 mice irradiated splenocytes for 7 days (1:1 ratio) Cells were labeled with immunofluorescent antibodies and analyzed for CD4+CD25+Foxp3-GFP+ve co-expression using flow cytometry.

CD4+ve cells incubated with anti-CD3 antibody in absence or presence of recombinant BTN2A2-Fc for day-1 and day-5 were used to extract RNA for qPCR analysis. Purified APCs (DC and B cells) from WT and BTN2A2-/- mice were co-cultured in suspension for 7 days with isolated murine T cells from Foxp3+ reporter mice for measurement of Tregs. CD4+ve T cells were isolated using EasySep™ Mouse CD4+ T Cell Isolation Kit (Stem Cell Tech); B cells using EasySep™ Mouse B Cell Isolation Kit (Stem Cell Tech); Pan-DC cells using EasySep™ Mouse Pan-DC Cell Enrichment Kit II (Stem Cell Tech); T cells using EasySep™ Mouse T Cell Isolation Kit (Stem Cell Tech) per protocols recommended by manufacturer.

To explore effects of BTN2A2 on Th17 differentiation, 1x10^6^ spleen cells were stimulated with soluble anti-CD3 antibody (0.5 μg/ml) and CD1 mice irradiated spleen cells (1:1 ratio) and Th17 differentiation cytokines cocktail (TGF-β1 1.5ng/ml, IL-6 10ng/ml, IL-1β 10ng/ml) with or without recombinant BTN2A2-Fc (10μg/ml) (Bettelli et al., 2006). Cells were co-cultured for 5 days and subjected to intracellular immunofluorescence staining of RORγt (Th17 cell marker). Cells were analyzed for CD4+RORγt +ve co-expression using flow cytometry. CD45 phosphatase inhibitor (Compound 211, Catalog # 530197; Millipore Sigma) that has previously been characterized as irreversible, and selective blocker of the allosteric pocket at the D1-D2 domains interface away from the substrate-binding/catalytic site (Perron et al., 2014), was used at 125nM concentration during the course of T cell differentiation studies.

### Flow Cytometry Analysis

Immune cells were freshly harvested from spleen and lymph nodes (inguinal, brachial and axillary) of mice. Cells were counted using a Hemavet 950FS hematology analyzer (Drew Scientific, Miami Lake, FL). Immunostaining and flow cytometry analysis of cells were performed as described previously(Li et al., 2018). Briefly, cells isolated from mice or treated in vitro cultures were harvested and stained with Ghost Dye™ Red 780 Viability Dye (Cell Signaling). Cells were washed extensively with 0.5% BSA in 1xPBS and resuspended in staining buffer (FBS); and further stained with fluorochrome conjugated antibodies for cells surface marker anti-CD3, anti-CD4 and anti-CD25. Cells were washed with staining buffer to remove excess antibodies and then fixed/permeabilized using mouse Foxp3 Buffer Set (BD Biosciences, San Jose, CA) according to manufacturer protocol. Fixed/permeabilized cells were incubated in rat serum for 15 minutes and stained with intracellular markers (Foxp3 and/or RORγt). Cells harvested from Foxp3-GFP transgenic mice were stained with anti-CD3, anti-CD4 and anti-CD25 antibodies, washed and suspended buffer containing DAPI. Cells were directly analyzed for CD4+CD25+Foxp3-GFP or CD4+RORγt expression. The stained cells were analyzed using a BD LSR Fortessa (BD Biosciences, San Jose, CA). Details of the commercial antibodies used in this study are provided in Supplementary Table 2.

For cell surface binding of BTN2A2-Fc, Jurkat cells were activated with anti-CD3/CD28 antibody (1μg/ml) for at least 48 hrs. Recombinant BTN2A2-Fc was conjugated with biotin in 1:2 ratio and pre-complexed with Strep-PE-Cy7 for hour on ice. Activated Jurkat cells (1x10^6^) were incubated with BTN2A2-Fc pre-complex (3μg/1x10^6^ cells) directly or preceding with anti-CD45RO (GeneTex Cat# GTX00596)/CD45RA (invitrogen Cat#MA1-19113) antibody (3μg/1x10^6^ cells) biding. Cells were washed with staining buffer and analysed in flow cytometer.

### Cells proliferation and detection of intracellular IL2

CD4+ cells were isolated from freshly harvested spleen of wild type B6 mice as described above. Cells were stained with CellTrace™ Violet dye at 5μM dye concentration in 10^6^ cells/ml dilution according to manufacturer protocol (Invitrogen).

2.5x10^6^ CellTrace™ Violet stained cells were incubated on bound anti-CD3 (0.5μg/ml) plus anti-CD28 (0.5μg/ml) with BTN2A2-Fc fusion protein (10μg/ml) or Fc control (10μg/ml). Cells were culture for 3 days at 37°C and 5% CO2 in RPMI medium supplemented with 10%FBS. Cell proliferation was analysed based on Dye dilution assay using flow cytometry analysis. For intracellular staining of IL2, cells were fixed, permeabilized and stained with anti-IL-2 antibody (Clone, JES6-5H4, Biolegend) and analyzed using a BD LSR Fortessa (BD Biosciences, San Jose, CA).

### Immunofluorescence Studies and Confocal Microscopy

Immunofluorescence staining for CD45 and CD3ζ was performed as described previously (Payne et al., 2020). Anti-CD3 antibody/anti-CD28 antibody (10μg/ml each) and recombinant BTN2A2-Fc (10μg/ml) were coated on glass cover slips in PBS overnight at 4°C. 1x10^6^ Jurkat cells pre-incubated with mouse anti-human CD45 (abcam cat# ab8216) in 75μl of 1xPBS for one hour were seeded on coated coverslips for 3 min at 37°C and fixed using Cytofix/Cytoperm-Fixation/Permeabilization Kit (BD biosciences). Coverslips were then gently washed two times with 1x PBS. Cells were blocked with blocking buffer (5% goat/mouse serum in 0.3% triton-x100 in 1xPBS) and stained with rabbit anti-human CD3ζ (abcam cat# Ab226263). Coverslips were then gently washed 3 time with 1x PBS for 10 minutes each and stained with Donkey anti-Mouse IgG Antibody, Alexa Fluor™ 555 and Goat anti-Rabbit IgG Antibody, Alexa Fluor™ 488 (ThermoFisher Scientific). Coverslips were then gently washed 3 times with 1x PBS, incubated in DAPI for 5 minutes and mounted on slides using ProLong™ Diamond Antifade Mountant (ThermoFisher Scientific). Immunofluorescence of stained cells was acquired with a Zeiss LSM 780 confocal microscope through a 63x resolution with oil immersion (refractive index 1.518). Orthogonal sections (z-stacks) of single cell were acquired using z-stack tool of confocal microscope. The image was analysed as an 8-bit image and intensity of red and green channel was measured from 0-255 grayscale fluorescent units. Co-localization analysis was performed on Image-J plugin “Colocalization Finder”. At least nine fields were analysed per experimental condition, in which an average of 500 T cells were analysed to yield the average amount of co- localization.

### Quantitative Polymerase Chain Reaction (QPCR)

For gene expression analysis, the total RNA was extracted using TRIzol™ Reagent (Thermo Fisher Scientific). The cDNA was synthesized from 500 ng of total RNA using SuperScript™ IV VILO™ Master Mix (Thermo Fisher Scientific). Quantitative real time PCR was performed with PowerUp™ SYBR™ Green Master Mix (Thermo Fisher Scientific) on a QuantStudio Real-Time PCR system (Thermo Fisher Scientific). GAPDH was used as internal controls. Gene expression was compared using the ΔΔCt method. Primers details are provided in (Supplementary Table 1).

### Animal Models

All animal experiments were conducted in accordance with the National Institutes of Health Guide for the Care and Use of Laboratory Animals and all the protocols used in this study were approved by the Cedars-Sinai Medical Center Institutional Animal Care and Use Committee.

Foxp3EGFP (C.Cg-*Foxp3^tm2Tch^*/J, Strain # 006769) mice: Foxp3EGF mice that co-express EGFP and the regulatory T cell-specific transcription factor Foxp3 under the control of the endogenous promoter was acquired from the Jackson Laboratory.

### BTN2A2 Knock-out mice (-/-)

BTN2A2<u>(-/-)</u> mice (strain: C57BL/6J-Btn2a2em1cyagen) were generated by deleting 10460 base pair region of btn2a2 gene comprising exon2-8 using CRISPR-Cas9 technique at commercial facility (Cyagen Biosciences) (Supplementary Figure 5A). Three PCR primers set (Supplementary Table 1) were used for genotyping mice using polymerase chain reaction method. Wildtype mice shows single band of 500bp, heterozygous shows two bands of 700bp and 500bp, while homozygous mice show 500bp single band on agarose gel electrophoresis (Supplementary Figure 5B).

### Glomerulonephritis Model

The detailed protocol to induce crescentic glomerulonephritis (GN) has been described previously (Saito et al., 2022). Nephrotoxic sera were raised in rabbits by repeated immunization with the purified glomeruli in complete and incomplete Freund’s adjuvant. The mice were preimmunized with normal rabbit IgG and complete Freund’s adjuvant five days prior to administration of nephrotoxic serum. Nephrotoxic serum nephritis was induced by the injection of 20 μl or 10 μl nephrotoxic serum intravenously at day 0. Four doses of 25 μg BTN2A2-Fc fusion protein or vehicle (control) were injected i.p. at day 0, 2, 4, 6 of nephrotoxic serum injection. Mice were scarified at day 7 to collect tissues, urine, blood cells and plasma. Immune cells were harvested from fresh whole spleen and lymph-nodes (Inguinal, Brachial and Axillary) and CD4+ve T cells were isolated using EasySep™ Mouse CD4+ T Cell Isolation Kit (STEM CELL TECH). Protein lysate of snap-freeze tissue in liquid nitrogen was prepared by homogenizing 5mg tissue in 500μl ice-cold RIPA lysis buffer. Lysate was agitated for 2 hrs at 4°C, centrifuged at 16000g for 20min at 4°C and supernatant was collected. About 80 μg of tissue lysate was used to detect IL17A expression using western blot.

To obtain the protein-to-creatinine ratio (mg/mg) in urine for evaluation of proteinuria, urine protein was measured with a protein assay dye (No. 500-0006, Bio-Rad, Hercules, CA), and urinary creatinine was measured using a Creatinine Assay Kit (No. DICT-500, BioAssay Systems, Hayward, CA). Fibrinoid necrosis (a precursor lesion for crescents, defined by finding GBM rupture, fibrin deposition, and karyorrhexis), and crescents were assessed in all glomeruli on one paraffin section for each mouse using periodic acid-methenamine silver and PAS stains, respectively. Representative images were captured using a light microscope (Nikon Eclipse 50i, Nikon, Tokyo, Japan). Histopathologic diagnosis of GN was evaluated by a board-certified renal pathologist (M.Y.)

### Miscarriage Model

DBA/2 and CBA/J mice were purchased from Jackson Laboratory. We crossed male DBA/2 with female CBA/J to document immunologically mediated pregnancy loss as described elsewhere (Zenclussen et al., 2005). As controls, we crossed female DBA/2 with male CBA/J strain and we evaluated litter size, pup weight and litter resorption rates. To study the therapeutic effects of BTN2A2, four doses of 25μg BTN2A2-Fc fusion protein were injected intraperitoneally at gestational days 6, 9. 12 and 15 in CBA/J female mice that were crossed with male DBA/2 mice. Animals were sacrificed at gestational day 18 and blood/tissues harvested for molecular analysis.

### Human Studies

This study was approved by the Institutional Review Board at the Cedars-Sinai Medical Center.

### Treg Assays

Whole blood was drawn from healthy unrelated individuals to prepare peripheral blood mononuclear cells (PBMC) using the Ficoll–Hypaque gradient centrifugation method. After the stimulator PBMC (unrelated) was irradiated, it was exposed to the responder PBMC at 1:1 ratio (1 × 10^6^/ml of each PBMCs) in the absence or presence of BTN2A2-Fc at 0, 10 μg/ml, and then incubated for 7 days for measurement of Tregs. MLR mixture were first stained with antibodies to CD45, CD3, CD4, CD25, and CD127. After permeabilization, the cells were stained with antibody to Foxp3. After acquiring cells by flow cytometry, lymphocytes separated from CD45+ leukocytes were plotted against CD4. CD4+ cells were plotted as CD25 versus CD127 and then CD25+CD127 ^low/−^ cells against Foxp3. CD4+/CD25+/CD127^low/−^/Foxp3+ cells were designated as Treg cells. Treg cell levels were expressed as Treg cell% in CD4+ T cells.

### Th17 assays

Human PBMCs (responder cells) isolated from Ficoll–Hypaque gradient centrifugation method were cultured in irradiated unrelated stimulator cells at 1:1 (1 × 10^6^/ml of each PBMCs) along with Th17 differentiation cytokines cocktail (anti-CD3 antibody 0.5μg/ml, TGFβ 1.5ng/ml, IL-6 10ng/ml, IL-1β 10ng/ml) with or without recombinant BTN2A2-Fc (10μg/ml) for 5 days. Cells were stained with surface marker CD3 and CD4. After permeabilization, the cells were stained anti-RORγt antibody and analyzed for CD4+RORγt +ve co-expression using flow cytometry. Th17 cells were expressed as RORγt +ve cells % in CD4 T cells. CD45 phosphatase inhibitor was used at 125nM concentration during the course of T cell differentiation studies.

### Statistics

Data were graphed and statistics performed using GraphPad Prism version 9.2. Data are presented mean ± standard deviation (SD) as indicated. Shapiro-Wilk test was used to test the normality of the data. Unpaired t-test was used to compare the data of two groups that passed normality test otherwise compared with Mann-Whitney test. For data with three or more groups, one-way ANOVA test with Tukey’s test for multiple comparison was used for data that passed normality test otherwise compared with Kruskal-Wallis test followed by Dunn’s test for multiple comparison. Statistically significant differences were defined as *p*<0.05 (*), *p*<0.01 (**), *p*<0.001 (***). The number of experiments per experiment is indicated in the legends to figures.

## Acknowledgements

We thank members of the Jordan and Karumanchi laboratory, and Nunzio Bottini for helpful discussions and insights. <u>Funding:</u> Internal Funding from Cedars-Sinai Medical Center. <u>Author Contributions:</u> Conceptualization: S.A., A.H.B., S.J., S.A.K; Methodology: S.A., R.Z., P.H., S.J; Cell Culture and Signaling Studies: S.A, R.Z., B.S., Animal models: M.Y., A.E., V.D; Structural Modeling and MST: R.M., M.K. Human Studies: R.Z., B.S., S.C.J., Writing, Reviewing and Editing: S.A., A.H.B., M.Y., S.Y., M.P., R.T., P.H., S.J., S.A.K.

## Data and Materials Availability

All data needed to evaluate the conclusions in the paper are present in the main manuscript or the Supplementary Materials. All reagents generated in this study including recombinant protein, baculoviral stocks, and knock-out mice will be made available to scientific community upon reasonable request.

## SUPPLEMENTARY MATERIALS

1. Supplementary Tables (Table 1: Primer Sequences; Table 2: List of Antibodies)

2. Supplementary Figures (1-5)

Supplementary Figure 1: Recombinant BTN2A2-Fc production

Supplementary Figure 2: BTN2A2-Fc signaling in Jurkat cells

Supplementary Figure 3: Co-immunoprecipitation studies for Zap70/CD45 in the presence of BTN2A2-Fc

Supplementary Figure 4: Expression of BTN2A2 and CD45 in T cells and Co- immunoprecipitation studies of Zap70/Cd43 in activated T cells treated with BTN2A2

Supplementary Figure 5: Knock-down and Site directed mutagenesis of CD45 Supplementary Figure 6: Lectin binding on Jurkat cells

Supplementary Figure 7: CD45 isoform expression in T cells

Supplementary Figure 8: qPCR studies for candidate molecules in Treg/Th17 pathway during treatment with BTN2A2

Supplementary Figure 9: BTN2A2 null mice generation/characterization and BTN2A2 expression in Antigen Presenting Cells

## Notes

### Competing Interest Statement

The authors have declared no competing interest.

