## Supplementary data for "Butyrophilin 2A2 promotes T cell immunoregulation by enhancing CD45 phosphatase activity within the immune synapse"

**Supplementary Table 1:** Primer sequence

| **Primer name** | **Primer Sequence** |
|---|---|
| BTN2A2-AgeI-1F | ATAT-ACCGGT-ATGGAACCAGCTGCTGCTC |
| BTN2A2-EcoRV-IEGRMD-711R | ATAT-GATATC-GTCCATACGTCCCTCGAT-GACAGTTTCCTTCTCCTGGCC |
| BTN-Xbal-F | ATAT-TCTAGA-ATGGAACCAGCTGCTGCTC |
| BTN-SacI-R | ATAT-GAGCTC-ACTCATTTACCCGGAGACAG |
| BTN-His-SacI-R | ATAT-GAGCTC-ACTCAGTGATGGTGATGGTGATGGACAGTTTCCTTCTCCTG |
| Mus-GAPDH-qF | ACCACAGTCCATGCCATCAC |
| Mus-GAPDH-qR | TCCACCACCCTGTTGCTGTA |
| Mus-ActinB-qF | CGCCACCAGTTCGCCATGGA |
| Mus-ActinB-qR | TACAGCCCGGGGAGCATCGT |
| Mus-Foxp3-F | CCTGGTTGTGAGAAGGTCTTCG |
| Mus-Foxp3-R | TGCTCCAGAGACTGCACCACTT |
| Mus-RORC-qF | CACGGCCCTGGTTCTCAT |
| Mus-RORC-qR | GCAGATGTTCCACTCTCCTCTTCT |
| cyBTNKOms-R1 | AAAGATGGGGAGAGAAAGTCAAGAG |
| cyBTNKOms-F1 | CCTACCTAGTTTTACCTGAGGTTC |
| cyBTNKOms-F2 | GGTCCATTGATATCCAAAGCCAAA |
| IL2RB-mus-qF | CTCAAGTGCCACATCCCAGATC |
| IL2RB-mus-qR | AGCACTTCCAGCGGAGAGATCT |
| IL21-mus-qF | GCCTCCTGATTAGACTTCGTCAC |
| IL21-mus-qR | CAGGCAAAAGCTGCATGCTCAC |
| TBX21-mus-qF | CCACCTGTTGTGGTCCAAGTTC |
| TBX21-mus-qR | CCACAAACATCCTGTAATGGCTTG |
| GATA3-mus-qF | CCTCTGGAGGAGGAACGCTAAT |
| GATA3-mus-qR | GTTTCGGGTCTGGATGCCTTCT |
| SMAD3-mus-qF | GCTTTGAGGCTGTCTACCAGCT |
| SMAD3-mus-qR | GTGAGGACCTTGTCAAGCCACT |
| CD5- mus-qF1 | AGAACCAGGTCTTCTGCCAAGG |
| CD5-mus-qR1 | TTGTGGGTGGAGGTGTCGTTCT |
| CD45cr3F | CACCGGTTTGCAGAATTACCACAA |
| CD45cr3R | AAACTTGTGGTAATTCTGCAAACC |
| CD45cr5F | CACCGTAATATTTACCCACCACCC |
| CD45cr5R | AAACGGGTGGTGGGTAAATATTAC |
| CD45-PCR-F | GGCAATAGTAAGGTGAGTAAGGA |
| CD45-PCR-R | TATTGACGCCGCTCAAGAACT |
| CD45-N419A.F | ACCTGGAATCCCCCTCAAAGATCATTTCATGCTTTTACCCTCTGTTATATA |
| CD45-N419A.R | TATATAACAGAGGGTAAAAGCATGAAATGATCTTTGAGGGGGATTCCAGGT |
| CD45-N468A.F | CCTACATCATTGCAAAAGTGCAACGTGCTGGAAGTGCTGCAATG |
| CD45-N468A.R | CATTGCAGCACTTCCAGCACGTTGCACTTTTGCAATGATGTAGG |

**Supplementary Table 2:** Antibody list

| **Antibody list** | | |
|---|---|---|
| **Vendor name** | **Catalog #** | **Name** |
| Novus Biologicals | NBP2-61717 | BTN2A2/Butyrophilin 2 Antibody (6C7D2) |
| R&D system | AF8645 | Human BTN2A2/Butyrophilin 2A2 Antibody |
| Cell Signaling Tech | 13917S | CD45 (Intracellular Domain) (D9M8I) XP® Rabbit mAb |
| Abcam | ab8216 | CD45 antibody [MEM-28] (ab8216) |
| ThermoFisher Scientific | MA1-19113 | CD45RA Monoclonal Antibody (MEM-56) |
| ThermoFisher Scientific | 5788-RBM30-P1 | CD45RB Recombinant Rabbit Monoclonal Antibody (PTPRC, 2877R) |
| ThermoFisher Scientific | MA1-19452 | CD45RO Monoclonal Antibody (UCHL1) |
| Genetex | GTX00596 | CD45RO antibody [UCHL1] |
| Abcam | ab28104 | Biotin Anti-CD43 antibody [MEM-59] |
| Cell Signaling Tech | 13838 | IL-17A (D1X7L) Rabbit mAb (Mouse Specific) |
| R&D system | MAB484-100 | Mouse CD3ε Antibody |
| BD BioScience | 566685 | Purifed NA/LE Mouse Anti-Human CD3 |
| BD BioScience | 553294 | Purified NA/LE Hamster Anti-Mouse CD28 |
| BD Bioscience | 555725 | Purified NA/LE Mouse Anti-Human CD28 |
| BD BioScience | 562163 | Purified NA/LE Rat Anti-Mouse CD3 Molecular Complex |
| Cell Signaling Tech | 7074S | rabbit IgG, HRP-linked Antibody |
| ThermoFisher Scientific | 10710C | Rat Serum |
| R&D system | AB-105-C | Normal Rabbit IgG Control |
| Sigma | I4506-10MG | IgG from human serum |
| R&D system | MAB004 | Mouse IgG2B Isotype Control |
| Sino-Biological | 10702-HNAH | IgG1 Fc Protein, Human, Recombinant (C103S) |
| Abcam | ab52959 | Recombinant Anti-CD3 epsilon antibody |
| Abcam | ab68235 | Recombinant Anti-CD3 zeta (phospho Y142) antibody |
| R&D system | MAB37041-SP | Human Lck Antibody |
| Cell Signaling Tech | 2714S | Lck (V49) Antibody |
| Millipore-Sigma | SAB4300118-100UG | phospho-LCK (pTyr$^{394}$) antibody produced in rabbit |
| Cell Signaling Tech | 70926S | Phospho-LYN (Tyr397)/LCK (Tyr394)/HCK (Tyr411)/BLK (Tyr389) (E5L3D) Rabbit |
| Cell Signaling Tech | 2705S | Zap-70 (99F2) Rabbit mAb |
| Cell Signaling Tech | 2717S | Phospho-Zap-70 (Tyr319)/Syk (Tyr352) (65E4) Rabbit mAb |
| Abcam | ab226263 | CD3 zeta antibody |
| Cell Signaling Tech | 2118 | GAPDH (14C10) Rabbit |
| Cell Signaling Tech | 7076S | Anti-mouse IgG, HRP-linked Antibody |
| ThermoFisher Scientific | A-31570 | Donkey anti-Mouse IgG Antibody, Alexa Fluor™ 555 |
| ThermoFisher Scientific | A-11008 | Goat anti-Rabbit IgG Antibody, Alexa Fluor™ 488 |
| **Antibody used in Flow cytometry analysis** | | |
| BD Bioscience | 553062 | FITC Hamster Anti-Mouse CD3e |
| BD Bioscience | 555332 | FITC Mouse Anti-Human CD3 Clone UCHT1 (also known as UCHT-1; UCHT 1) (RUO) |
| BD Bioscience | 555349 | APC Mouse Anti-Human CD4 Clone RPA-T4 (RUO) |
| BD Bioscience | 553051 | APC Rat Anti-Mouse CD4 |

| | | |
|---|---|---|
| Biolegend | 102049 | Brilliant Violet 711™ anti-mouse CD25 Antibody |
| BD BioScience | 562894 | BV421 Mouse Anti-Mouse RORγ |
| Biolegend | 100206 | PE anti-mouse CD3 Antibody |
| BD Bioscience | 566684 | PE Mouse Anti-Human CD3 Clone OKT3 (RUO) |
| BD Bioscience | 560046 | PE Mouse anti-Human FoxP3 Clone 259D/C7 (RUO) |
| BD BioScience | 563081 | PE Mouse anti-Human RORγt |
| BD BioScience | 562607 | PE Mouse anti-Mouse RORγt |
| BD BioScience | 555275 | PE Rat Anti-Mouse CD3 Molecular Complex |
| BD Bioscience | 560408 | PE Rat anti-Mouse Foxp3 |
| Biolegend | 503808 | PE anti-mouse IL-2 Antibody |
| BD Bioscience | 553022 | PE Rat Anti-Mouse CD5 |
| BD BioScience | 564219 | Human BD Fc Block |
| BD BioScience | 553141 | Purified Rat Anti-Mouse CD16/CD32 (Mouse BD Fc Block™) |
| Biolegend | 405206 | Strep-PE-Cy7 |
| BD BioScience | 562868 | BV421 Rat IgG1, κ Isotype Control |
| BD BioScience | 554648 | PE Mouse IgG2a, κ Isotype Control |
| BD Biosciences | 555848 | PE Rat IgG2b, κ Isotype Control |

A

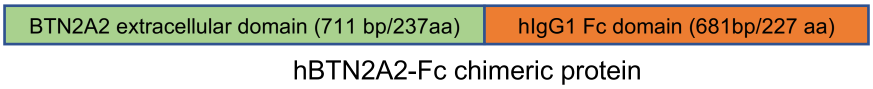

B

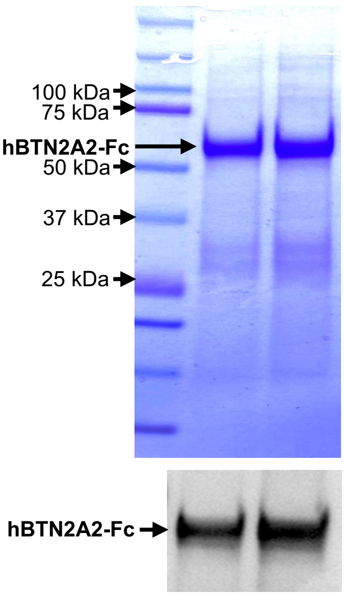

C

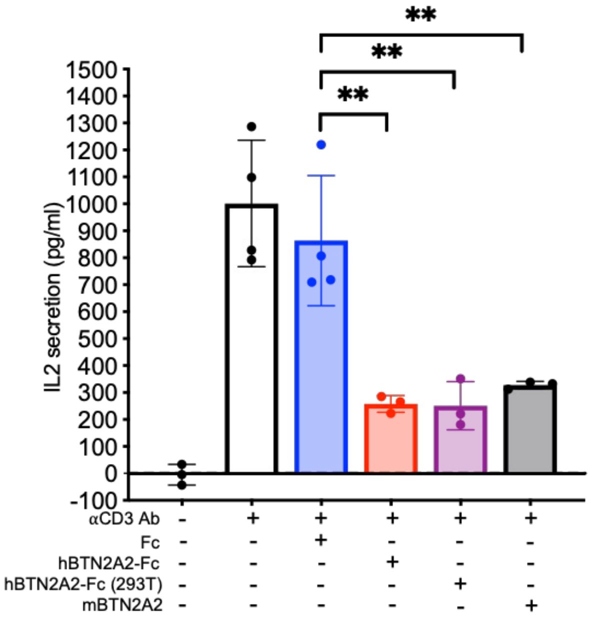

**Supplementary Fig 1:**

A. Schematic diagram showing the recombinant human BTN2A2-Fc fusion protein. The recombinant fusion gene consisting of the two extracellular domains (IgV and IgC2) of BTN2A2 molecule (green color) cloned along with the Fc region of a human IgG1 antibody (orange color). The chimeric protein (BTN2A2-Fc) is separated by IEGRMD spacer polypeptide.

B. Coomassie blue staining of purified recombinant BTN2A2-Fc protein on reducing (SDS) gel with a protein ladder in the left column and a band corresponding to BTN2A2-Fc (~55 kDa) in the right two columns. Lower panel shows western blot of recombinant BTN2A2-Fc using specific anti-human BTN2A2 antibody.

C. IL-2 secretion from Jurkat cells incubated for 24hrs with plate-immobilized anti-CD3 antibody (1μg/ml) in the absence or presence of immobilized recombinant BTN2A2-Fc (10μg/ml), 293-cell derived human BTN2A2-Fc (R&D systems) or mouse BTN2A2 (R&D systems), and immobilized immunoglobulin Fc domain as negative control is depicted (N=3 or 4).

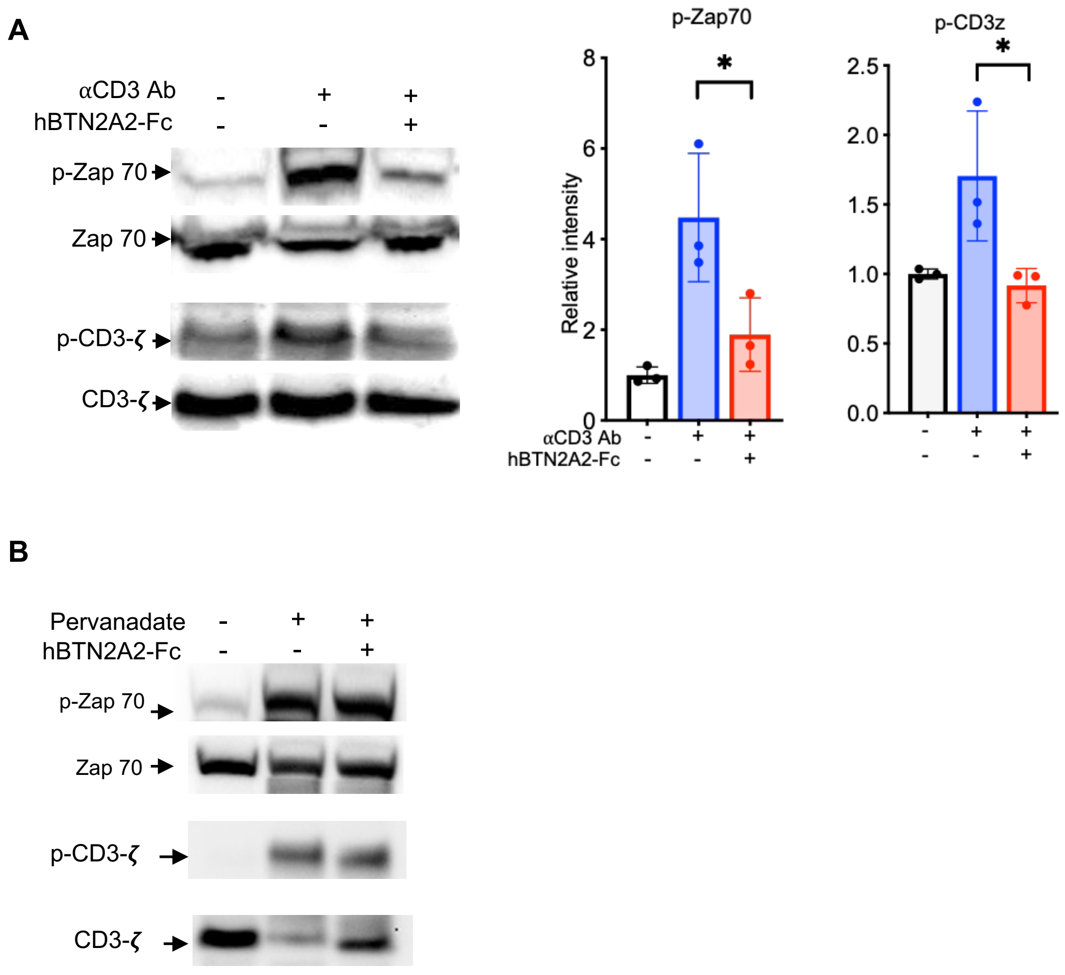

**Supplementary Fig 2:**

A. Immunoblot analysis of phosphorylated proteins, p-Zap70 and p-CD3-ζ and total Zap70 and CD3-ζ in Jurkat cells stimulated for 3 min with immobilized anti-CD3 antibody (10 µg/ml) in the presence of immobilized recombinant BTN2A2-Fc (10 µg/ml) or immunoglobulin Fc domain (10 µg/ml). Right panel shows relative intensity plots as mean±SD (N=3).

   One-way ANOVA with Tukey's multiple comparison test; $p<0.05$ (*), $p<0.01$ (**), $p<0.001$ (***).

B. Jurkat cells were treated with permanent phosphatase inhibitor (Pervanadate) plus minus BTN2A2-Fc or left untreated. After cell lysis, phosphorylated proteins, p-Zap70 and p-CD3-ζ were analyzed by Western blot. Total Zap70 and CD3-ζ were used as loading controls.

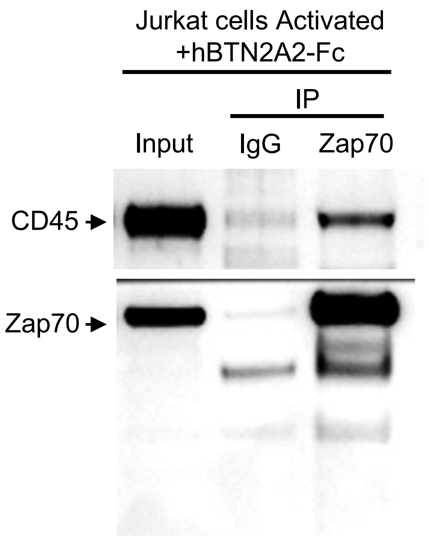

**Supplementary Fig 3:** Jurkat cells were stimulated for 3 min with immobilized anti-CD3 antibody (10 μg/ml) in the presence of BTN2A2-Fc (10 μg/ml). Cells were lysed in IP buffer, immunoprecipitated with anti-Zap70 antibody and immunoblotted for CD45.

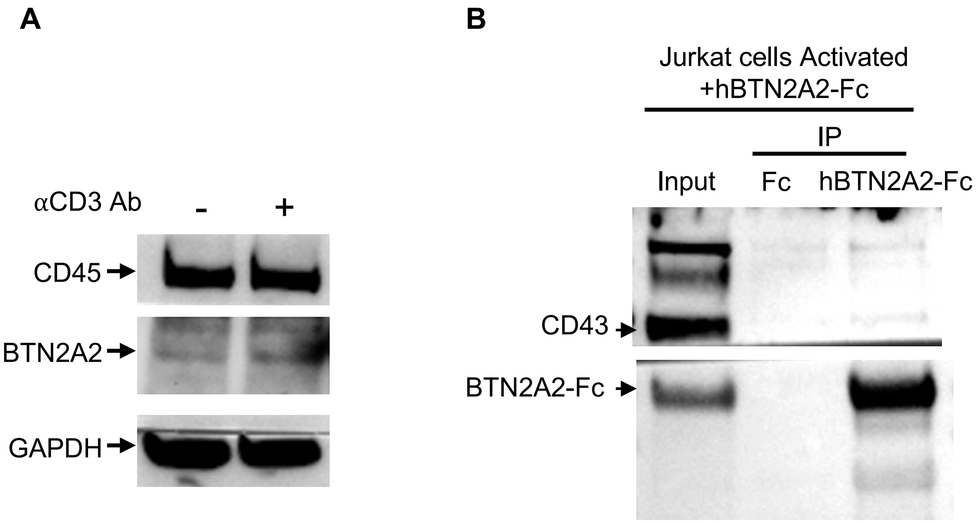

**Supplementary Fig 4:**

A. Jurkat cells were stimulated with/without anti-CD3 antibody (1μg/ml) for 48 hrs and the lysates were immunoblotted with anti-CD45 or anti-BTN2A2 antibody. GAPDH expression was used as a loading control.
B. Jurkat cells were treated with anti-CD3 antibody and anti-CD28 antibody (1μg/ml each) for 48 hours and lysed in IP buffer. Co-immunoprecipitation of cell lysate with recombinant BTN2A2-Fc or Fc-tag protein overnight and protein G. Immunoblotting performed with anti-CD43 antibody. Input is ~2% of cell lysate.

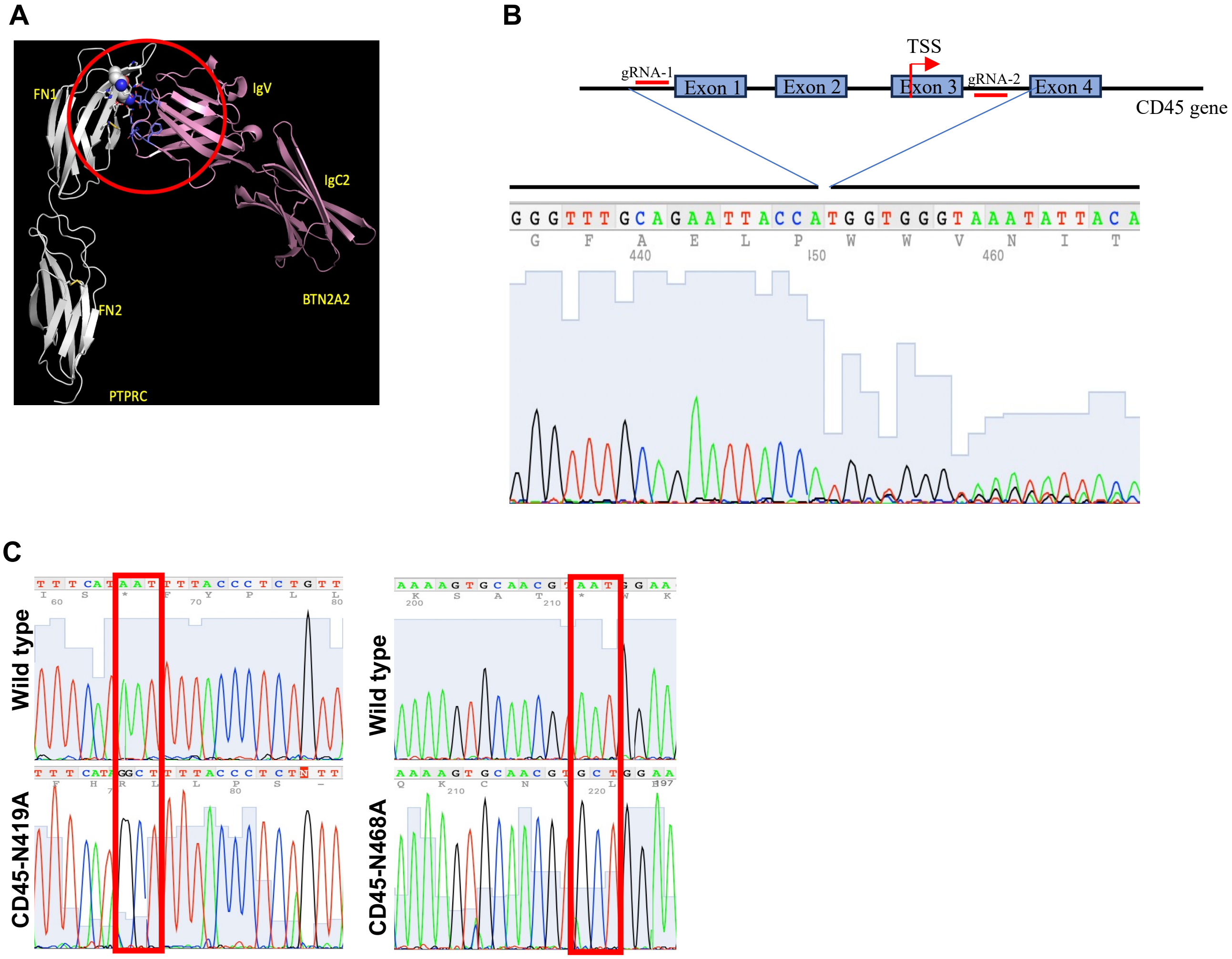

**Supplementary Fig 5:**

A. Ribbon diagram representation of protein-protein interaction between extracellular domains of BTN2A2 (pink) and CD45 (PTPRC)(white) showing potential amino acids (blue sticks) determining their interaction.
B. Schematic of CD45 gene showing location of gRNA sequences complementarity. Lower panel show electropherogram of the sequence deleted upstream of exon-1 to downstream of exon-3.
C. Electropherogram showing change of nucleotides, AAT to GCT at Asn419Ala and AAT to GCT at Asn468Ala in CD45 mutants gene.

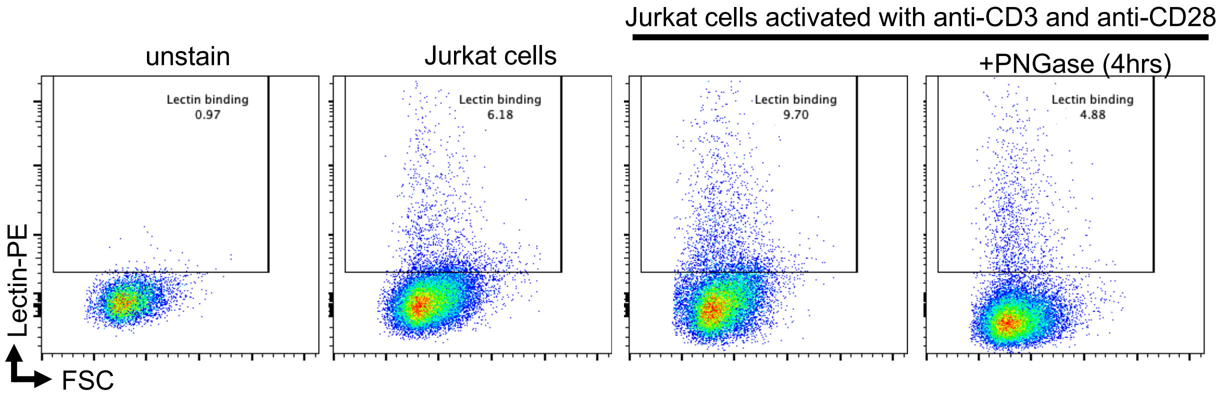

**Supplementary Fig 6:** Lectin binding on Jurkat cells stimulated with and without anti-CD3 and anti-CD28 antibody and treated with PNGase enzyme.

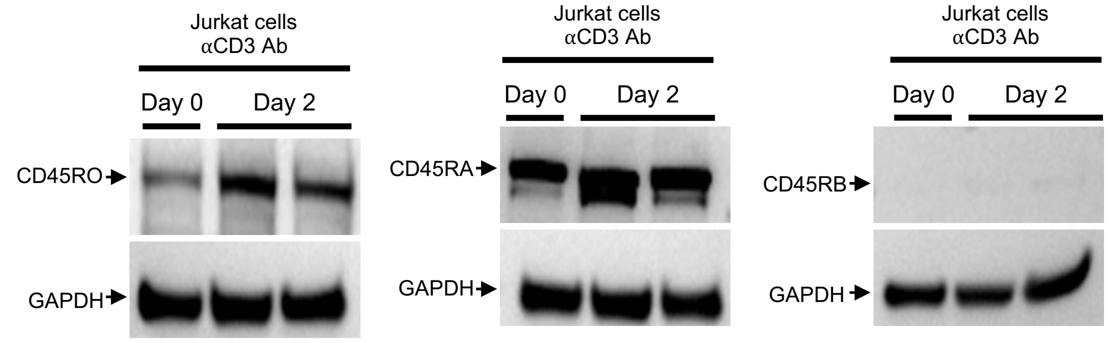

**Supplementary Fig 7:** Western blot demonstrating expression of the CD45 isoforms at baseline and at 48 hours after stimulation with anti-CD3 antibody in Jurkat cells.

Data are depicted as mean ± SD (n=4). Unpaired t-test; $p<0.001$ (***).

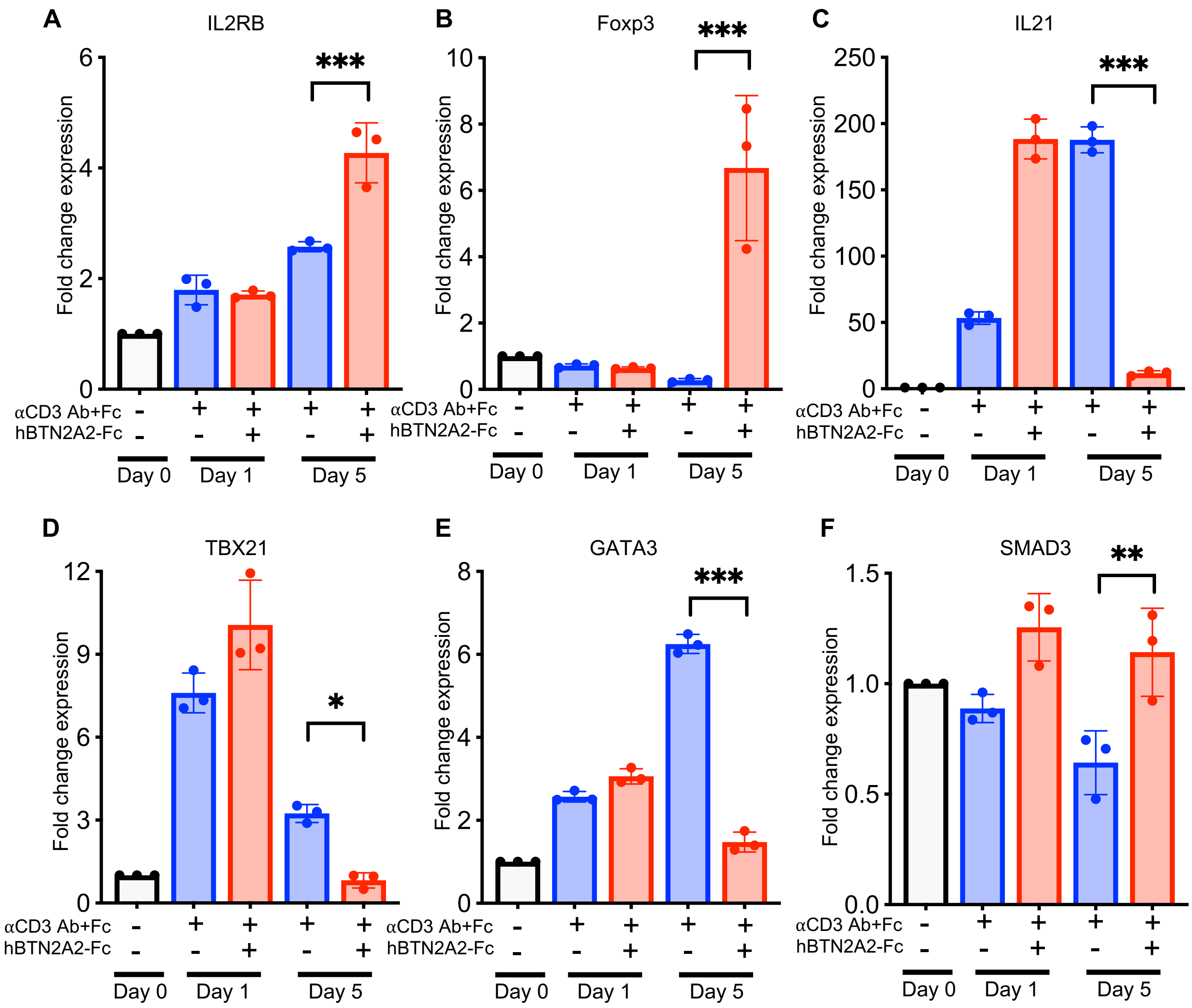

**Supplementary Fig 8:** Quantitative PCR analysis for IL-2RB (A), Foxp3 (B), IL-21 (C), TBX21(D), GATA3 (E) and Smad-3 (F) in primary CD4 cells incubated for 1 Day and 5 days in absence or presence of anti-CD3 antibody and/or recombinant BTN2A2-Fc. All data represented as mean ± standard deviation. n=3 per group for all experiments. One-way ANOVA with Tukey's multiple comparisons test; $p$<0.05 (*), $p$<0.01 (**), $p$<0.001 (***).

**A.** Overview of the Targeting Strategy

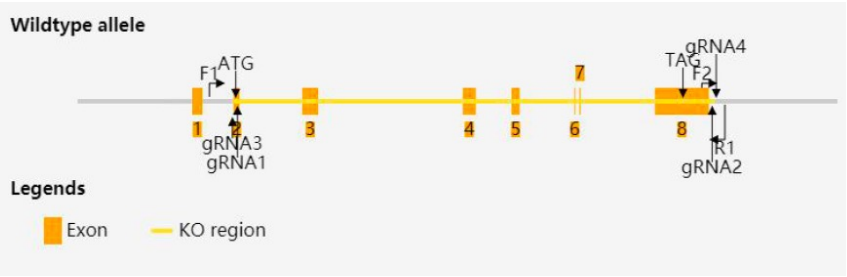

**gRNA target sequence**

```
gRNA1 (matching reverse strand of gene): AGGCTATCGGGCAAGAACGCAGG
gRNA2 (matching reverse strand of gene): AAATCCGATTCATGTGGACCTGG
gRNA3 (matching forward strand of gene): GTCATGTACATGAGAACCGTGGG
gRNA4 (matching reverse strand of gene): TAGTGAGACCTATCTTGAAGGGG
```

**Genotyping Primer sequence**

```
Forward primer (F1): 5'-CCTACCTAGTTTTACCTGAGGTTC-3'
Forward primer (F2): 5'-GGTCCATTGATATCCAAAGCCAAA-3'
Reverse primer (R1): 5'-AAAGATGGGGAGAGAAAGTCAAGAG-3'
```

**B.** Mouse genotyping Results

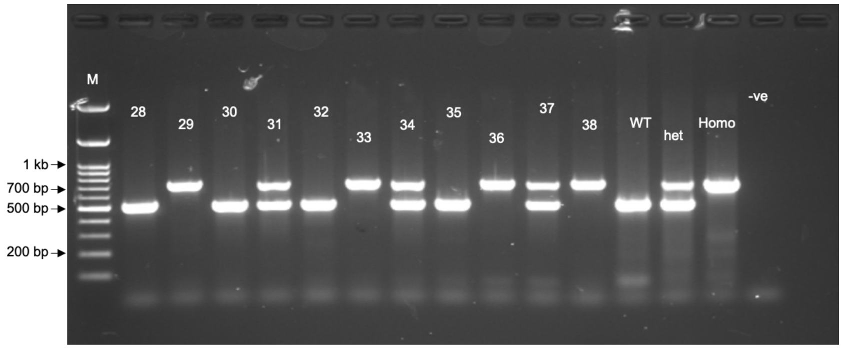

**C**

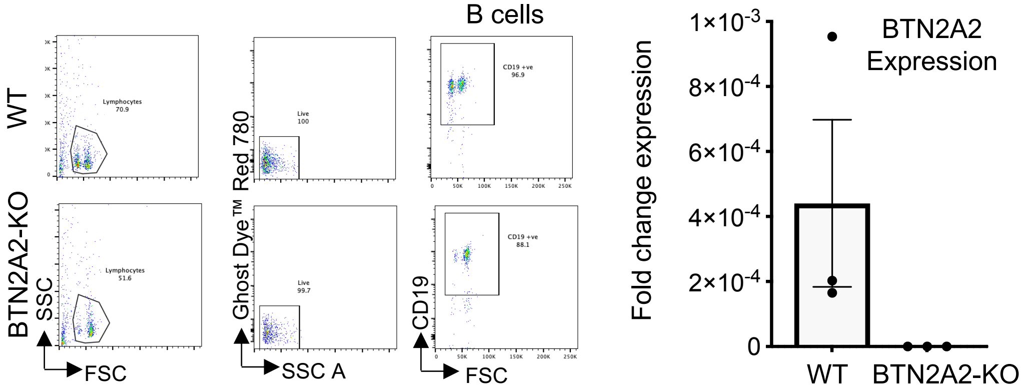

**D**

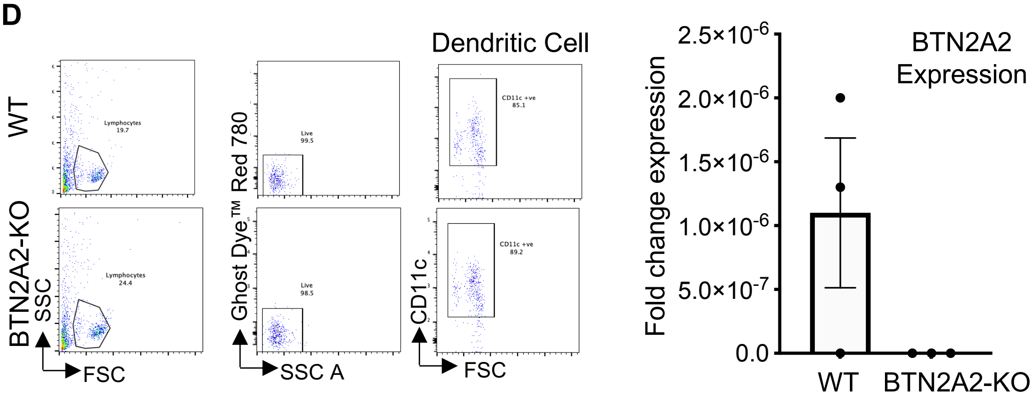

**Supplementary Fig 9:**

**Supplementary Fig 9:**

A. Btn2a2$^{-/-}$ mouse model (C57BL/6J) was generated by CRISPR/Cas-mediated genome engineering (cyagen.com). The Btn2a2 gene (NCBI Reference Sequence: NM_175938; Ensembl: ENSMUSG00000053216) is located on mouse chromosome 13. Eight exons are identified, with the ATG start codon in exon-2 and the TAG stop codon in exon-8 (Transcript Btn2a2-203: ENSMUST00000110433). Exons 2~8 (covers 100.0% of the coding region) were selected as target site. Cas9 and gRNA co-injected into fertilized eggs for KO mouse production. The pups were genotyped by PCR followed by sequencing analysis. The size of effective KO region: ~10500 bp.

B. Agarose gel electrophoresis of genotyping PCR. Homozygotes (Homo) mouse shows one band with 700 bp, heterozygotes (Het) mouse with two bands at 700 bp and 500 bp; wildtype mouse (WT) one band at 500 bp.

C-D qPCR analysis of BTN2A2 expression in B cells (C) and Dendritic cells (D) populations purified using magnetic beads from spleen and lymph nodes of wildtype and BTN2A2-KO mice. Left panels shows flow cytometry plots of purified cells, and right panels shows fold change in RNA expression of BTN2A2 transcript compared to GAPDH.
